# Identification of *cis*-Acting Elements that Regulate Influenza A Virus Segment 7 Differential mRNA Splicing

**DOI:** 10.64898/2025.12.22.695654

**Authors:** Shu Zhou, Rute M. Pinto, Hou Wei Chook, Hui Min Lee, Eleanor R. Gaunt, Paul Digard

## Abstract

Influenza A virus segment 7 encodes the M1 matrix protein and M2 ion channel from an unspliced primary transcript (mRNA1) and spliced mRNA2, respectively. Both proteins are essential for efficient virus replication, but the intrinsic sequences that determine segment 7 splicing balance are not fully defined. Here, we systematically investigated how conserved *cis*-acting splicing elements regulate segment splicing. We found that while the splice acceptor sequence is optimal, both potential upstream polypyrimidine tracts regulate splicing, with the 5’ region acting positively and the 3’ region negatively to maintain M1 expression. Nucleotide A713 is the most likely mRNA2 branch point, but potentially with redundancy from other local adenosines. Mutagenesis of the mRNA2 splice donor sequence showed that it is nearly optimal but that nucleotides 51 and 55 modestly downregulate splicing efficiency. Most introduced mutations that change splicing balance are nonsynonymous in M1. Nevertheless, polypyrimidine tract and branch point mutations that significantly perturbed M1:M2 balance resulted in viable viruses. In contrast, the splice donor sequence was largely intolerant of mutation. We conclude that M1:M2 balance is maintained via suboptimal splice donor and polypyrimidine tract splicing signals, with the latter allowing more evolutionary flexibility to vary the ratio and/or M1 sequence.

## Introduction

Influenza A virus (IAV) is an enveloped negative-sense single-stranded RNA virus with eight genome segments. Ten proteins encoded by these segments are essential for virus fitness outside of the laboratory: segments 1-6 encode polymerase basic subunit 1 (PB1), polymerase basic subunit 2 (PB2), polymerase acidic subunit (PA), haemagglutinin (HA), nucleoprotein (NP), and neuraminidase (NA). Segments 7 and 8 encode more than one protein by alternative splicing: matrix protein (M1) and matrix 2 ion channel protein (M2) from segment 7, and nonstructural protein 1 (NS1) and nuclear export protein (NEP) from segment 8 [1]. The viral proteome is further elaborated through a variety of transcriptional and translational mechanisms to produce a strain-dependent suite of accessory protein species [2].

The M1 protein has multiple functions. As well as forming a helical filament under the virus envelope, it determines viral particle morphology [3,4] and recruits other viral components to assemble at the site of budding [5,6]. It also binds to viral ribonucleoproteins (RNPs) in a regulated manner to control their nuclear import and export [7]. The M2 protein is a proton channel that is translated from the main spliced transcript of segment 7. It mediates acidification of the interior of the virus particle in endosomes during virus entry [8,9] and has also been proposed to drive membrane scission during virus budding [10]. Although both M1 and M2 proteins are found in the virion, the copy number of M2 is two orders of magnitude lower than of M1 [11]. Nevertheless, expression levels are more equal in cells, with M2 comprising between 10% to 60% of segment 7 derived protein production, depending on the cell and virus strain [12], implying that M1 and M2 expression is regulated. Primarily, this is thought to occur at the level of mRNA splicing. Newly transcribed segment 7 mRNA molecules accumulate in nuclear speckles for splicing from the earlier stages of infection and although measurements of the ratio of mRNA2:mRNA1 varies according to time-point, virus strain, host cell and assay method, the unspliced transcript tends to predominate [12-15].

Segment 7 mRNA splicing is more complex than simply producing mRNA2. At least four spliced transcripts have been identified [16-19] in addition to unspliced mRNA1. The spliced mRNAs use different splice donor (SD) sequences but share the same splice acceptor site (SA). Spliced mRNA2 encodes the M2 ion channel protein [8]. Spliced mRNA3 is not required for virus replication *in vitro* and no encoded protein has yet been identified [20]. mRNA4 encodes an M2 variant (M42) with a distinct ectodomain but identical C-terminal domain [21]. All IAV strains produce unspliced mRNA1 and mRNA2, but the abundance of mRNA3 is variable among strains and only a small minority of viruses produce mRNA4 [18,21].

Four *cis*-acting elements are required for cellular pre-mRNA splicing: an SD at the 5’- end of the intron/exon boundary; and three separate but usually juxtaposed elements: a branch point (BP) and polypyrimidine tract (PPT) at the 3’end of the intron and a 3’-SA sequence overlapping the intron/exon boundary (Figure 1A). These motifs are recognised by the spliceosome, based on their match to the conserved consensus sequences [22-25]. Most introns have highly conserved termini: a GU dinucleotide at the 5’end and an AG dinucleotide at the 3’ end [26-28]. These lie within broader consensus nucleotide sequences for SD and SA sites, which are MAG|GURAGU and YAG| respectively [29-31]. The BP consensus sequence (YUN<u>A</u>Y, with the underlined A being the actual branch point) is more loosely conserved, while the PPT is usually a stretch that contains around 10 pyrimidines located 5-40 nucleotides upstream of the SA site [32]. The SD sequence is recognised by spliceosome component U1 snRNP through RNA-RNA base pairing to initiate splicing [29,33]. After cleaving the transcript at the 5’ end of the intron, an intronic guanine at the 5’-end conjugates to the BP adenine to form a lariat structure [32]. The spliceosome’s two U2 auxiliary factor subunits (U2AF65 and U2AF35) recognise the PPT and SA, respectively, recruiting U2 snRNP and promoting spliceosome assembly [25,34]. Finally, the two exons ligate and the intron in lariat configuration is excised and degraded [32].

**Figure 1.**
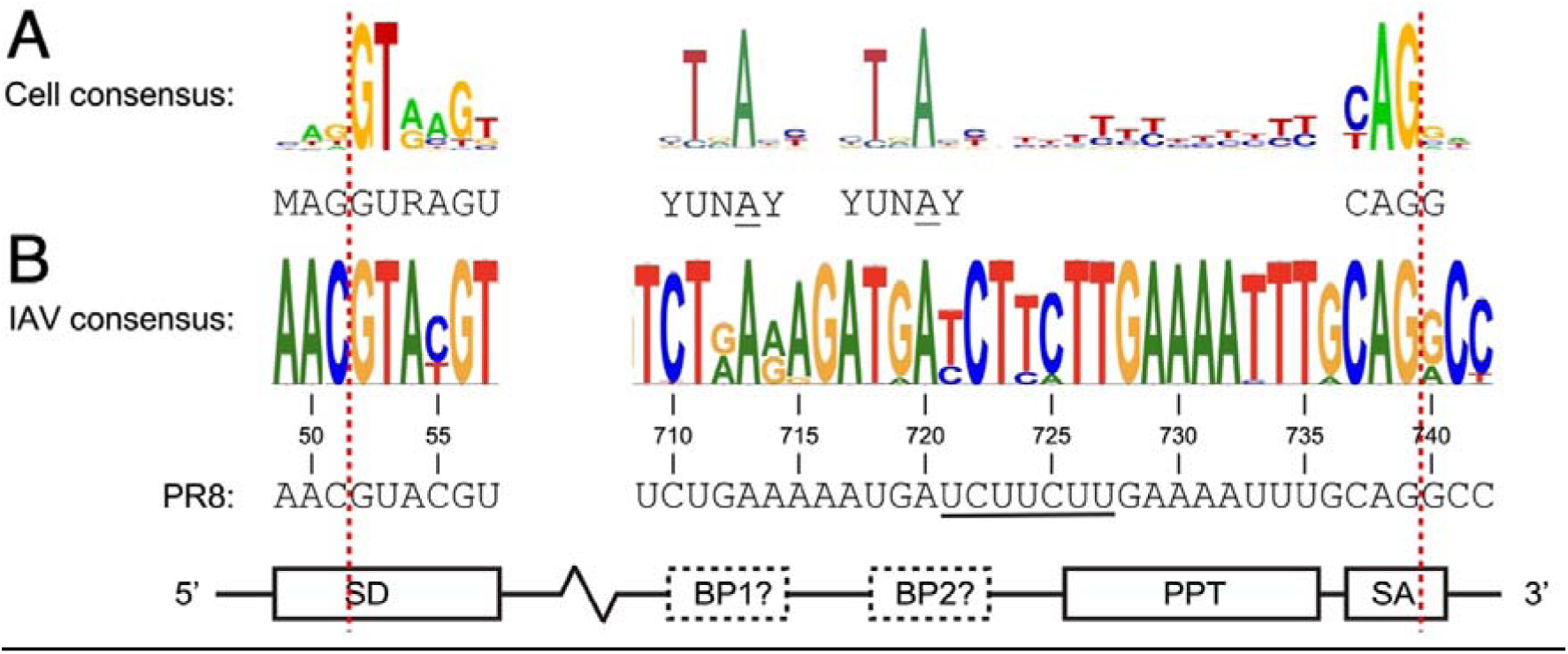
Consensus alignments of cellular and IAV segment 7 *cis*-acting splice elements. Sequence logos for (A) a selection of experimentally validated mammalian mRNAs (SD, PPT and SA motifs, adapted with permission from [47,48]; BP motifs adapted from [49] under CC BY 4.0 permission) and (B) IAV segment 7 sequences are shown, along with the cellular consensus (M indicates A or C and Y indicates C or U/T). The sequence of the A/Puerto Rico/8/34 (PR8) virus strain is also shown. Note that the sequence logos were generated from sequences downloaded from NCBI and thus follow the convention of T rather than U for RNA. Red dashed lines indicate splice sites (boundary between intron and exon). Boxes indicate splice site elements. SD: Splice donor; BP: Branch point; PPT: Polypyrimidine Tract (following the cellular consensus location and length - the underlined region from nucleotides 721-727 indicates an alternative prediction [46]); SA: Splice Acceptor. Dashed boxes indicate that the location of IAV segment 7 BP is unclear.

Since the cellular spliceosome mediates IAV segment 7 mRNA splicing [35-37], the core splicing signals in IAV mRNAs generally, though imperfectly, resemble those of host cell transcripts. In contrast to most cellular pre-mRNAs, both unspliced and spliced transcripts of segment 7 are translated to essential viral proteins. Therefore, IAV must only splice a fraction of segment 7 transcripts to ensure correctly balanced M1 and M2 production. One possibility as to how this is achieved is that viral factors ensure limited splicing, and there is evidence to suggest that segment 7 splicing is regulated by NS1 and/or the viral polymerase complex [35-39]. However, segment 7 mRNA is incompletely spliced even when expressed by cellular RNA polymerase II (rather than viral polymerase) in the absence of other IAV proteins [39-41], indicating that viral proteins are not the only regulatory factors determining splicing efficiency. For cellular mRNAs, auxiliary splicing regulatory elements (SRE), including exonic and intronic splicing enhancers and silencers (ESE, ESS, ISE, ISS) can also play a role in modulating splicing efficiency [42,43]. Similarly, segment 7 also contains SREs, and various cellular proteins such as NS1-BP, hnRNP K and ASF/SF2 have been proposed to to modulate mRNA2 production [35,36,38,39,44]. However, at least some of these interactions appear to be strain- and/or host-specific. For example, the identity of segment 7 nucleotide (nt) 334 can determine whether TRA2A is recruited to inhibit splicing, but only a minority of viruses possess a consensus target sequence [45]. Similarly, the region surrounding the SA (nt 707-779) is reported to modulate the splicing ratio in certain virus strains in a cell type-dependent manner [19]. Furthermore, the majority of reported interactions are proposed to increase rather than decrease segment 7 splicing. Thus, for a virus with a notoriously broad host range, the relative contribution that SREs and the core-splice site sequences make to ensure incomplete splicing of IAV segment 7 remain unclear. The segment 7 mRNA2 splice sites are imperfect matches for the core cellular consensus sequences (Figure 1) and based on mutational studies of cellular splicing, these mismatches are likely to impair the recognition of *trans*-acting elements [31], which may result in reduced splicing efficiency. However, splice site efficiency is difficult to predict from sequence [31]. Other uncertainties are that the segment 7 BP has never been experimentally mapped, only computationally predicted, while the extent and location of the PPT also remain unclear [46].

In this study, we set out to test the hypothesis that the primary balance between M1 and M2 mRNAs is set by the conserved *cis*-acting splice site elements of IAV segment 7, by systematically characterising the strength and location of the SD, BP, PPT and SA motifs. We find that segment 7 has a strong SA and potentially redundant BP sequences, and that the key suboptimal splicing signals that give the desired mRNA1/M1 and mRNA2/M2 ratios are the SD and PPT signals.

## Materials and Methods

### Bioinformatics analyses

To generate a consensus sequence for IAV segment 7 mRNA2 splice sites, 13,280 full-length IAV segment 7 sequences (all subtypes and host species) were downloaded from the National Centre for Biotechnology Information (NCBI) server. For some analyses, sequence metadata was used to split the dataset according to host species. A script was written to extract the host information from the sequence metadata. Host details were then manually reviewed, and a second script was used to further group sequences because of variability in nomenclature and in numbers of individual sequences returned. Specifically, because of the diversity of avian hosts, sequences isolated from birds were grouped into a larger ‘avian’ category. Similarly, sequences isolated from swine, pig, *sus scrofa* or wild boar were grouped into a ‘swine’ category, those from equine, Equus ferus caballus, donkey or horse were grouped into an ‘equine’ category, canine, *Canis lupus familiaris*, *Vulpes vulpes*, or dog samples were grouped into a ‘canine’ category and feline or tiger samples were grouped into the ‘feline’ category. Sequences that did not contain a host species identifier or which contained human or *homo sapiens* were assigned to the ‘human’ dataset. Sequences labelled as reassortant viruses were excluded. Viral sequences were aligned and logos were generated using WebLogo [47,48].

Counterpart sequence logos representing the consensus SD, SA and PPT sequences from experimentally verified human genes, originally abstracted from the Exon-Intron database [50], were adapted with permission from https://weblogo.berkeley.edu/examples.html. The consensus BP sequence from the bovine genome was modified with permission (https://creativecommons.org/licenses/by/4.0/) from reference [49]. To assess the strength of potential splice site motifs in segment 7, the A/Puerto Rico/8/1934 (PR8) segment 7 sequence (GeneBank accession: EF467824) was truncated at the polyadenylation signal (to mimic the mRNA sequence) and used as input for published prediction programmes: Human Splicing Finder (HSF) [51], MaxEntScan [52] and SVM-BPfinder [53]. HSF was run through the web portal (https://genomnis.com/hsf), using HSF scoring. For SVM-BP, the Python script (https://github.com/comprna/SVM-BPfinder) was run on a high-performance Linux Research Compute Cluster at the University of Edinburgh. The maxlen parameter was increased to 1000 to scan the entire segment. For MaxEntScan, the Python script (https://github.com/kepbod/maxentpy) was run on Google Colab (https://colab.research.google.com/). SD inputs used 9-mers of the GT dinucleotide flanked by three upstream and four downstream bases, while SA inputs were 23-mers of the AG dinucleotides preceded by 18 upstream and three downstream residues. SD and SA strengths were calculated by score5 and score3 functions. Differences relative to WT (ΔMaxEnt score) were visualised as a heatmap using the seaborn package (Python). A diverging colour scale centred at zero was used to represent decreases or increases in predicted SD or SA strength.

Local minimum free energy (MFE) RNA secondary structures of WT and mutant PR8 segment 7 sequences spanning the SA site (nt 707-769), corresponding to the sequence used in previous studies [46,54] were predicted using RNAfold. Local RNA accessibility was predicted using RNAplfold (version 2.4.17). Both programmes were run using the ViennaRNA package [55], with previously described parameters (-W 200, -L 150 and -u 20 for RNAPplfold) [56]. Accessibility was estimated as the probability that a continuous nucleotide window of a set length remained unpaired. Depending on the analysis, probabilities for windows of 4, 7, and 15 nt were extracted from the RNAplfold output and plotted using Matplotlib (Python) and Prism 11. Window lengths were selected to encompass the 7-nt predicted PPT or 15-nt PPT identified in this study and the extended SA (CAG|G) motif, respectively, as calculating accessibility over a functional motif has been shown to perform better than assessing the accessibility of single nucleotides [57].

### Cells, viruses and plasmids

Madin-Darby canine kidney (MDCK) cells, human lung A549 cells (ATCC) and human embryonic kidney (HEK 293T) cells (ATCC) were cultured according to standard procedures [58]. Cells were routinely tested for mycoplasma contamination. PR8 [59] and A/California/04/2009 (H1N1) (Cal04) [60] wild type (WT) and modified viruses were rescued by transfection of eight plasmid systems containing the viral genome segments under bidirectional polI and polII promoters into HEK 293T cells using Lipofectamine 2000 following manufacturer’s instructions. Rescued viruses were propagated in MDCK cells as previously described [58]. Site-directed mutagenesis of segment 7 was performed using Q5 Site-Directed Mutagenesis Kits (New England Biolabs, E0554) following the manufacturer’s instructions. All constructs were confirmed by Sanger sequencing (GENEWIZ). To titre viruses, confluent monolayers of MDCK cells were infected with serially-diluted viral samples in the presence of viral growth medium (serum free DMEM containing 0.14% BSA and 1 µg/ml TPCK trypsin) and incubated for one hour, followed by addition of 1.2% Avicel (Dupont) overlay at 1:1 (v/v) for PR8 or 1:2 for Cal04. After 48 (PR8) or 72 (Cal04) hours, plates were formalin fixed and stained with 0.1% toluidine blue (Sigma). To quantify plaque size, fixed plates were scanned in an Epson Perfection V750 Pro Scanner at a resolution of 360 dpi. Images were converted to grey scale, contrast was adjusted and the area of each plaque was determined in ImageJ 2. 5-20 plaques were selected and measured per experiment and normalised to the values from WT virus. For protein and mRNA analysis in single-cycle infections, cells were infected at an MOI of 5, incubated for one hour and then cells were washed to remove input viruses and medium was replaced.

### Virion RNA extraction and sequencing

Viral genomic RNA was extracted from 140 µl of MDCK-grown virus stocks using a QIAamp Viral RNA Kit (QIAGEN) and the addition of 5.6 µg of carrier RNA according to the manufacturer’s instructions, then DNase-treated using TURBO DNA-free kits (Thermo Fisher Scientific). Reverse transcription was performed using Superscript IV (Thermo Fisher Scientific) and a Uni 12 primer [61] that targets genome segment termini (AGCAAAAGCAGG). 1 µl of the resulting cDNA was used for PCR using PrimeSTAR GXL Premix (Takara Bio Inc). Forward and reverse primers AGCAAAAGCAGGTAGATATTG and AGTAGAAACAAGGTAGTTTTTTAC respectively were used to generate a segment 7 amplicon. PCR conditions were 98°C for 5 min, 35 cycles of 98°C for 10s, 55°C for 15s and 68°C for 2 min. PCR products were purified by Monarch DNA Gel Extraction Kit (New England Biolabs) and analysed by Sanger sequencing (GENEWIZ).

### Minireplicon assay

HEK 293T or A549 cells were seeded in 24-well plates at 2 x 10^5^ cells per well and transfected the next day with PR8 or Cal04 reverse genetics plasmids expressing combinations of PB2, PB1, PA, NP, NS and M segments of each strain. For HEK 293T cells, 50 ng of each plasmid (total of 300 ng DNA per transfection) was transfected using 1 µl lipofectamine 2000 (Thermo Fisher Scientific), whereas for A549 cells, 100 ng of each plasmid (total 600 ng DNA per transfection) was transfected using 2 µl lipofectamine 2000 following the manufacturer’s instructions. The empty vector plasmid containing bi-directional RNA PolI and PolII promoters (“pDUAL”) [59] was used as a negative control by replacing the segment 7 plasmid. Medium was removed 48 hours after transfection, and cells were lysed using Laemmli buffer (20% glycerol, 2% Sodium Dodecyl Sulfate (SDS), 100 mM Dithiothreitol (DTT), 24 mM Tris pH 6.8, 0.016% bromophenol blue) for western blotting or lysed for total RNA extraction as described below.

### RT-PCR and splicing analysis

Total RNA was extracted from virus-infected or plasmid-transfected cells using a RNeasy Mini Kit (QIAGEN) according to the manufacturer’s instructions. After DNAse treatment using a TURBO DNA-free Kit (Thermo Fisher Scientific), cDNA was generated from 800 ng of RNA using Lunascript RT Master Mix Kit (New England Biolabs) and oligo dT (IDT) as a primer. The cDNA was PCR amplified using PrimeStar Max Premix (Takara Bio Inc.). M segment PCR was performed with the following primers: forward primer: GAGTCTTCTAACCGAGGTC for constructs with SD mutations or TTCTAACCGAGGTCGAAAC for the rest of the analyses and a common reverse primer: CCACAGCACTCTGCTGTTCCT. mRNA1 and mRNA2 were distinguished based on the sizes of their PCR products: 939 or 934 bp for mRNA1 and 251 or 246 bp for mRNA2, depending on the forward primer used. NP mRNA PCR was performed with forward primer: ACGGAAAGTGGATGAGAGAACTCAT and reverse primer: CAACCGACCCTCTCAATATGAGTGC. PCR conditions were 98 °C for 5 min, 23 (for segment 5) or 28-30 cycles (for segment 7, depending on experiment) of 98 °C for 10s, 55 °C for 15s and 68 °C for 10s. PCR products were separated in 2% agarose gels stained with SYBR™ Safe DNA Gel Stain (Thermo Fisher) and imaged using an Odyssey Fc imaging system (Li-Cor Inc.). PCR products were analysed by Sanger sequencing (GENEWIZ) and Unipro UGENE [62].

### Western blotting analysis

SDS-PAGE followed by western blotting was performed using home-made 20% or 4-20% gradient gels (BioRad) and the Trans-Blot Transfer System (BioRad) according to the manufacturer’s instructions. Nitrocellulose membranes with 0.2 µm pores were blocked with phosphate-buffered saline (PBS)/0.1% Tween20/5% skimmed milk at room temperature for one hour, washed with PBS/0.1% Tween20 and incubated with primary antibody diluted in blocking buffer. Following overnight incubation at 4 °C, membranes were washed with PBS/0.1% Tween20 and incubated with fluorophore-labelled secondary antibody diluted in blocking buffer for one hour, washed and then imaged using an Odyssey Fc imaging system and Image Studio Lite software (Li-Cor Inc.). Purchased monoclonal antibodies were against α-tubulin (Thermo Fisher Scientific, Clone YL1/2), IAV M2 (Abcam, Clone 14C2), and IAV NP (Abcam, Clone C43). Purchased polyclonal antibodies were against IAV M1 (GeneTex, GTX125928) and M2 (Thermo Fisher Scientific, PA5-32233). In-house polyclonal rabbit anti-NP (antiserum A2915) and anti-NS1 (antiserum V29) have been previously described [63]. Secondary antibodies were IRDye 800CW anti-Mouse (Li-Cor Inc, 925-32212), IRDye 680RD anti-Mouse (Li-Cor Inc, 926-68070), IRDye 680RD anti-Rat (Li-Cor Inc, 926-68076) and IRDye 800CW anti-Rabbit (Li-Cor Inc, 926-32213). Densitometric analysis of western blots was done using the gel plot function in ImageJ 2 [64].

### Statistical Analyses

Statistical analyses of numerical data were conducted from at least 3 biologically independent repeats. Graphical data were plotted and analysed using GraphPad Prism 11. Statistical tests used an ordinary One-way ANOVA test. Multiple comparisons were performed against WT using Dunnett’s post-test. **** = P<0.0001; *** = P<0.001; ** = P<0.01; * = P<0.05; absence of significance symbols = P>0.05. Error bars indicate the standard deviation (protein expression analysis) or standard error of the mean (plaque size comparisons).

## Results

### Bioinformatic analysis of segment 7 splicing *cis*-acting elements

Splicing of cellular mRNAs is directed by well-defined but degenerate consensus sequences [31]. Outside of the almost obligatory GU and AG nucleotides at either end of the intron, individual pre-mRNA sequences are rarely complete matches to the consensus (Figure 1A) and this can affect splice site selection [65]. The SD sequence of IAV segment 7 mRNA2 is a reasonable match to the cellular consensus, possessing the obligatory intron boundary dinucleotide as well as matching at nt 49, 50, 54, 56 and 57 (Figure 1B). Consistent with this, both MaxEnt and HSF splice site prediction programmes scored the PR8 SD sequence highly (9.51 versus cellular median values in the 7-10 range [52,66,67] and 80.98 where a score > 80 is categorised as strong respectively [51]). The SA site is an even better match to consensus; perfect in many strains and varying only at position 740 in a minority of isolates (Figure 1B). The viral PPT has been suggested to be composed of a stretch of 7 pyrimidines running from position 721 - 727 [46], but this would be both shorter and spaced further away from the SA than the norm for cellular transcripts [68]. The region immediately 5’ to the SA site where cellular PPTs are usually found is not especially pyrimidine-rich, having a run of 5 purines upstream of three uracils (Figure 1B). MaxEnt and HSF both consider SA and PPT sequences as one unit and returned scores for PR8 segment 7 that fall below the cellular median value (MaxEnt: 6.30), or into the ‘less strong’ category (HSF: 78.95), consistent with a weaker PPT component. A conserved adenosine at position 720 has been predicted to be the segment 7 BP [46], based on its contextual match to the cellular consensus. However, A713 is in a similar context, only deviating from consensus at the 3’-end of the motif. The SVM-BP programme [53] designed to identify BP sites scored A713 as slightly more likely to be the branch point than A720 (0.06 vs -0.22 respectively, while HSF scored A713 highly (83 compared to a validated dataset average of 83.4) and failed to identify A720 as a potential BP. Thus overall, visual and computational inspection of the segment 7 splice site motifs suggest varying degrees of match to the cellular consensus, with the PPT sequence in particular looking atypical.

These sequence analyses provided predictions of splice site strength, but did not establish the functional contribution of each *cis*-acting element. We therefore experimentally tested the importance of the predicted splice signals by systematic mutagenesis, starting from the 3’-end SA site. To avoid confounding effects from non-synonymous mutations in the M1 gene or changes to overlapping segment-specific RNA packaging signals [58] while retaining expression via the viral polymerase, we utilised a transfection-based minireplicon system, using segment 7 as the reporter construct [37]. To further mimic viral infection, we included segment 8, given the potential effects of NS1 on segment 7 mRNA splicing, nuclear export and overall cellular translation [13,37,69-72].

### PR8 segment 7 has an optimal SA sequence

The core sequence of the PR8 segment 7 SA is an exact match to cellular consensus. However, to test for context-dependent effects, we examined the consequences of mutating the nucleotides either side of the essential AG motif, as not all cellular SA sites meet the extended consensus (Figure 1) [73,74]. Accordingly, two mutants were made in which the AG motif of the SA was preceded by either A or G at position 737 instead of C (Figure 2A). Uracil (U) was not tested as this would introduce a premature stop codon into the M1 gene and was therefore not a viable evolutionary strategy for the virus. To estimate how the introduced mutations were expected to affect SA intrinsic strength, we used MaxEntScan [52], which predicted that both C737A and C737G mutations would weaken the SA. We then tested segment 7 RNA synthesis and protein expression from the mutated segments using the minireplicon system, measuring mRNA levels and relative M2:M1 ratios by RT-PCR and western blotting respectively. The WT segment produced a mix of unspliced mRNA1 and spliced mRNA2 (Figure 2B) as expected [37,38], as well as appreciable quantities of both M1 and M2 polypeptides (Figure 2C). However, the C737A and C737G mutations (immediately upstream of the AG dinucleotide) strongly inhibited both mRNA splicing and M2 expression, reducing the latter to below the level of detection when replicate experiments were quantified (Figure 2D). Therefore, 737C is a key component of the segment 7 SA, consistent with its 100% conservation in segment 7 sequences (Figure 1) [75] and the likelihood that this nucleotide will cause reduced splicing in cellular mRNA [74].

**Figure 2.**
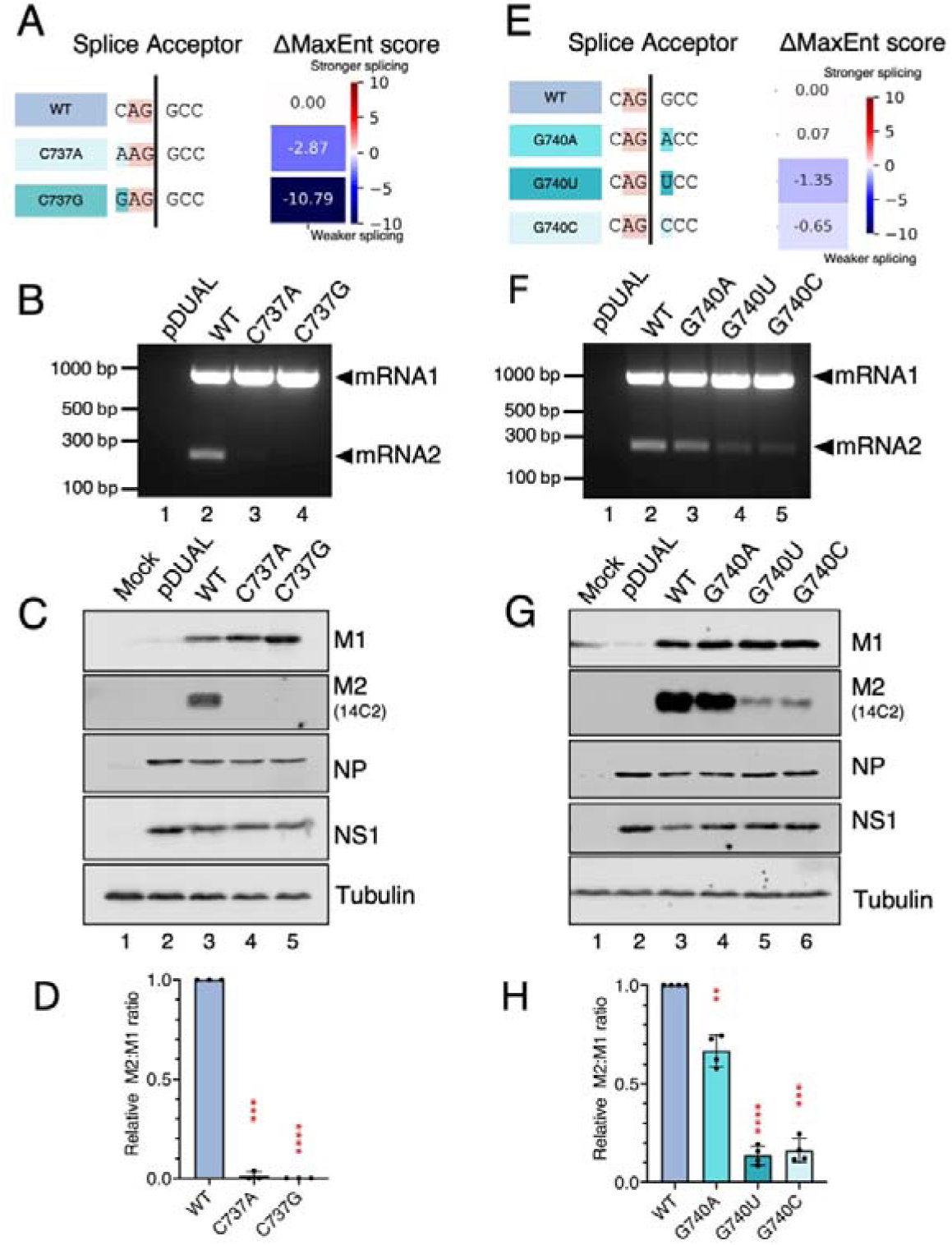
The IAV segment 7 SA has a close to optimal sequence. (A and E) Left - design of mutations (blue shades) to change the nucleotide preceding or following the acceptor AG dinucleotide (orange). Right - heatmap showing predicted ΔMaxEnt (mutant – WT) scores for indicated mutants. Positive values indicate the mutation is predicted to increased SA strength, whereas negative values indicate a weaker SA. (B-D and F-H) HEK 293T cells were co-transfected (or mock transfected) with the viral polymerase and NP (3PNP) and NS segment plasmids along with the indicated WT or mutant segment 7 constructs (pDUAL indicates an empty vector control without segment 7) and harvested 48 h later. (B, F) Total RNA was extracted from cells and segment 7 mRNAs detected using RT-PCR followed by agarose gel electrophoresis. (C, G) Cells were lysed in Laemmli buffer and protein expression was analysed by SDS-PAGE and western blotting. (D, H) Relative M2:M1 ratios were calculated by densitometry from at least three independent experiments. Statistical annotations are the result of an ordinary One-way ANOVA with Dunnett’s post-test relative to WT. **** = P<0.0001; *** = P<0.001; ** = P<0.01; * = P<0.05; absence of significance symbols = P>0.05. Error bars indicate the standard deviation.

In contrast to nucleotide 737, naturally occurring single nucleotide polymorphisms (SNP) are seen at segment 7 nt 740 (after the AG dinucleotide), albeit with a clear preference for G and A over C or U (Figure 1). MaxEntScan predicted that substitutions to C or U would reduce the strength of SA markedly, whereas A was predicted to have only a minor effect (Figure 2E). To test this, we made the corresponding constructs (Figure 2E) and monitored their splicing efficiency by RT-PCR and western blotting as before. WT segment 7 exhibited the highest splicing efficiency by the qualitative RT-PCR assay (Figure 2F), while all 740 mutants yielded reduced amounts of the M2 protein in correlation with the amount of spliced transcript (Figures 2G, H). However, the G740A variant was less inhibitory than the other two mutants (Figure 2F-H). Thus, the G/A SNP at position 740 may contribute to regulation of segment 7 splicing in some strains of IAV [19,76].

Previous studies by Moss *et al*. and Bogdanow *et al.* suggested that alterations in RNA conformation surrounding IAV segment 7 SA can influence splicing by modulating its accessibility to *trans*-acting factors [19,46,54]. Therefore, to assess whether our mutations might be affecting splicing indirectly by inducing structural changes that influence spliceosome recognition, we compared *in silico* mRNA structures of WT and SA motif mutants using RNAfold. The local secondary structure prediction of WT segment 7 (Supplementary Figure 1A) was a bulged-hairpin with the A of the AG dinucleotide at the end of a stem and the splice site in the unpaired region, similar to that reported by previous studies [19,46,54]. Both C737A and C737G disrupted the base-pairing status of the AG dinucleotide, pushing the splice site further into a more open bulge (Supplementary Figure 1B). To evaluate whether this change was reflected across all potential RNA structural ensembles, local RNA structural accessibility of the WT and mutant SAs were analysed using RNAplfold (Supplementary Figure 1C). The predicted accessibility of the extended SA motif (CAG|G) increased in both C737A and C737G mutants, suggesting that the mutations are unlikely to reduce spliceosome recognition by decreasing local RNA accessibility. In contrast, G740U and G740C substitutions extended the base-paired stem downstream of the AG dinucleotide and reduced the size of the loop containing the SA (Supplementary Figure 1D), whereas the G740A mutation had a relatively minor effect on the overall predicted structure (Supplementary Figure 1D). Analysis of local RNA structural accessibility further indicated that the probability of the extended SA motif being unpaired was substantially reduced in G740 mutants compared with WT (Supplementary Figure 1E), providing a plausible explanation for the reduced splicing efficiency observed experimentally. Potentially therefore, the C737 and G740 mutants downregulated splicing through partially distinct mechanisms: C737 substitutions by directly disrupting sequence-specific recognition by U2AF35, while G740 substitutions may influence both U2AF35 recognition and alter local RNA structure. Irrespective of mechanism, none of the tested mutations in the segment 7 SA enhanced splicing efficiency, suggesting that IAV segment 7 SA is inherently strong.

### The segment 7 polypyrimidine tract is key to setting M1:M2 balance

PPT sequences function via recruiting spliceosomal U2AF, which then brings the U2 snRNP to the BP which is required for lariat formation [25,34]. The PPT is located upstream of the SA [32] and its strength is determined by proximity and continuity: a long PPT near to the SA is generally stronger than a short one or one further from the SA [77]. The PPT of segment 7 has been predicted to be nucleotides 721-727 of the pre-mRNA (underlined in Figure 1) [46], but this would be both short and relatively far from the SA. Alternatively, a more conventionally sited PPT ending two nucleotides upstream of the SA at position 735 would extend 5’-wards into the predicted PPT but is interrupted by 5 consecutive purines centred at position 730. To experimentally test these options, we first mutated pyrimidines at nucleotide positions 721 through 727 to either adenines or guanosines whilst avoiding the creation of cryptic SA (AG) motifs (Figure 3A, mutants 721-24A and 724-27G). MaxEntScan predicted that both sets of purine substitutions would weaken the intrinsic strength of the overall 3’-splice site. As before, pre-mRNA splicing in the minireplicon system was measured using RT-PCR and western blotting for M1 and M2. Supporting the identification of the 721-727 region as a PPT, mRNA2 splicing and M2 protein production were decreased when pyrimidines were replaced by purines (Figure 3B lanes 3-4 and C lanes 4-5, Figure 3D for quantification). However, the two sets of mutations affected splicing differentially. While mRNA2 splicing was still detected for the 721-24A mutant, 724-27G almost totally abrogated splicing and subsequent M2 expression.

**Figure 3.**
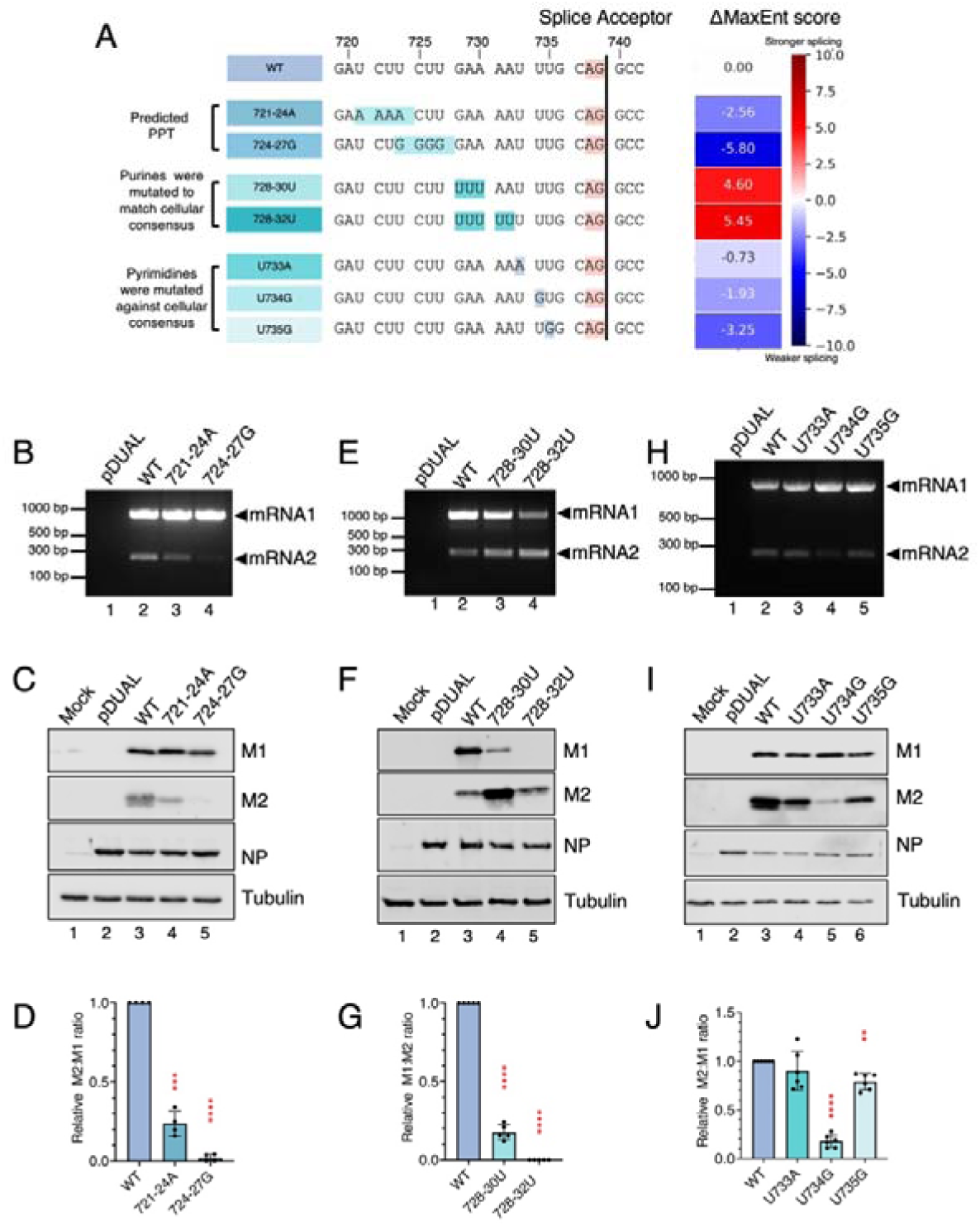
Identification of the segment 7 PPT. (A) Left - sequences of segment 7 mutants with altered PPTs. Mutations are highlighted in blue shades and the segment 7 SA is highlighted in orange. Right - heatmap showing predicted ΔMaxEnt (mutant – WT) scores for indicated SA mutants. Positive values indicate the mutation is predicted to increased SA strength, whereas negative values indicate a weaker SA. (B-J) The RNP components, NS segment and the indicated segment 7 or pDUAL empty vector plasmids were co-transfected (or mock transfected) in HEK 293T cells. (B, E, H) Segment 7 mRNA was detected by RT-PCR followed by agarose gel electrophoresis. (C, F, I) Protein expression was analysed by SDS-PAGE and western blotting. (D, G, J) Densitometric quantification of the relative (D, J) M2:M1 ratios and (G) M1:M2 ratio was analysed by an ordinary One-way ANOVA test with Dunnett’s post-test relative to WT (N = 4-6). **** = P<0.0001; *** = P<0.001; ** = P<0.01; * = P<0.05; absence of significance symbols = P>0.05. Error bars indicate the standard deviation. The ratio in (G) is reversed to avoid divide by zero errors.

While the effect of mutations within nucleotides 721-727 confirmed that segment 7 sequences up to position 721 play a role in PPT function, they did not necessarily define its full extent. Therefore, we mutated regions downstream of the predicted PPT to either match (mutants 728-30U and 728-32U) or violate (U733A, U734G and U735G) the cellular PPT consensus. MaxEntScan predicted that the increased pyrimidine content would strengthen the intrinsic 3′ splice-site strength, whereas purine replacements would weaken it. RT-PCR results from these constructs showed that mRNA1 levels decreased and mRNA2 levels increased with the introduction of uridines (Figure 3E, lanes 3-4), while removing uridines at positions 734 and 735 had the converse effect (Figure 3H, lanes 4 and 5), consistent with uridine content close to the SA site modulating splicing. Further supporting this, all up and down mutations also caused significant alterations to the balance of M1 and M2 protein expression (Figures 3F, G, I and J; note that the y axis of G is reversed relative to panels D and J). However, while reduced mRNA1 levels in the 728-32U mutant correlated with a drastically lowered M1:M2 ratio, overall segment 7 protein expression was low (Figure 3F, lane 5). This particular mutation introduced a 10-nucleotide long polyuridine stretch, which in other contexts has been shown to induce viral polymerase slippage [78-80], potentially providing an explanation for reduced levels of M1. In agreement, Sanger sequencing of PCR products spanning the region showed a marked reduction in sequence fidelity at this region (data not shown).

As described above, we next assessed whether changes in local RNA folding could contribute to the effects of PPT mutations on segment 7 splicing. Compared with WT RNA, the 728–730U mutation (which increased splicing) increased predicted accessibility of both the extended 15 nt PPT region identified in this study and the previously proposed 7 nt PPT (Supplementary Figure 2A-D). In contrast, all other PPT mutations were predicted to either slightly increase or have little effect on the local accessibility of both PPT regions despite reducing splicing efficiencies. We also investigated whether the effects of PPT mutations could be explained by altered accessibility of the SA motif. Almost all PPT mutations were predicted to increase accessibility around the CAG|G motif, with the exception of 728–730U (Supplementary Figure 2E and F). However, these predicted changes did not correlate with the observed reduced splicing efficiency. Thus, overall, while changes in local RNA structure may contribute to the phenotype of mutations that increased segment 7 splicing, decreased U2AF65-mediated PPT recognition because of reduced pyrimidine content remains a plausible explanation for the observed splicing defects.

Overall, these data indicate that segment 7 contains a bipartite PPT spanning nucleotides 721-735, comprising stronger 5’ and weaker 3’ elements, with the latter acting to limit oversplicing and thereby preserve M1 expression. RNA secondary structure predictions suggest that sequence-dependent PPT recognition, rather than RNA accessibility, is the dominant factor regulating splicing in this region, potentially mediated by U2AF65.

### Segment 7 has potentially redundant branch points

Splicing of cellular pre-mRNAs requires the formation of a lariat structure [32]. The BP is an essential *cis*-acting element lying 5’ to the PPT that interacts with the SD site to initiate lariat formation. In most cases, the BP is an adenine residue, as its 2’-OH group is necessary to attack the phosphate backbone of the SD sequence [32]. Although they are essential, directly identifying BP(s) can be difficult because of lariat degradation [81]. However according to the scanning model for 3′ splice-site recognition, AG dinucleotides located within approximately 12–15 nt of the BP are bypassed during splice-site selection, whereas AG dinucleotides located further downstream of the BP, if present, would compete with the canonical SA and therefore are normally selected against – thus defining an ‘AG exclusion zone’ or AGEZ [53,74,82]. Consequently, the AGEZ provides a constraint on the likely location of the BP(s). In PR8 segment 7, the first AG upstream of the canonical SA is located at nt 701-702. Combining this with the PPT location (nts 721-735) inferred from our previous data (Figure 3) narrows the likely BP(s) location to within the range of nts 703-720. However, unlike cellular pre-mRNAs, the influenza A virus genome is highly compact and individual RNA sequences are often constrained by multiple overlapping functions.

Consequently, the absence of upstream AG dinucleotides may not solely reflect splice-site selection. To experimentally map the AGEZ, our strategy was to insert CAG triplets (the C to enhance splicing efficiency and therefore experimental sensitivity) at various positions upstream of the canonical splice acceptor (Supplementary Figure 3A). All CAG triplets were designed to be in frame with the downstream M2 ORF if they were used for splicing. SA usage was then investigated by minireplicon, RT-PCR and Sanger sequencing. Unspliced mRNA1 and spliced mRNA2 were detected from WT segment 7 as expected (Supplementary Figure 3B lane 8). Splicing of the 725CAG construct was blocked (lane 6), while all other mutants spliced to some extent. Sanger sequencing of the spliced mRNA PCR products demonstrated that all mutants with a CAG triplet upstream of nucleotide 725 still utilised the canonical SA (Supplementary Figure 3C). However, the novel SA created at nucleotide 728-730 (728CAG) was used in preference to the WT CAG. These data indicate that the functional AGEZ, defined as the region where introduction of a novel AG interferes with use of the canonical SA, begins no later than nucleotide 725 (Supplementary Figure 3A). Therefore if the segment 7 BP follows the scanning model for 3′ splice-site recognition [74], the most likely branch-point region is upstream of nt 714.

Previous bioinformatic analyses predicted nucleotide 720 as the segment 7 BP [46], but this position falls within the AGEZ region identified by our mapping, when it would be more conventional for the BP to be upstream of the AGEZ. To investigate the previously predicted BP experimentally, we mutated A720 to each of the other three nucleotides (Figure 4A; BP2). Reanalysis of the mutant sequences using MaxEntScan predicted slight increases in splicing (Figure 4A, ΔMaxEnt score). If nucleotide 720 were the sole BP, these mutations would be expected to destroy its function, block splicing and reduce M2 protein expression. However, no reduction in M2 expression was detected for any of the three mutants, which instead gave slightly increased (but not statistically significant) M2:M1 ratios relative to WT segment 7 (Figure 4B and C). Thus, either nucleotide 720 is not the segment 7 BP or an alternative BP can be utilised in its absence, consistent with its proximity to the segment 7 AGEZ.

**Figure 4.**
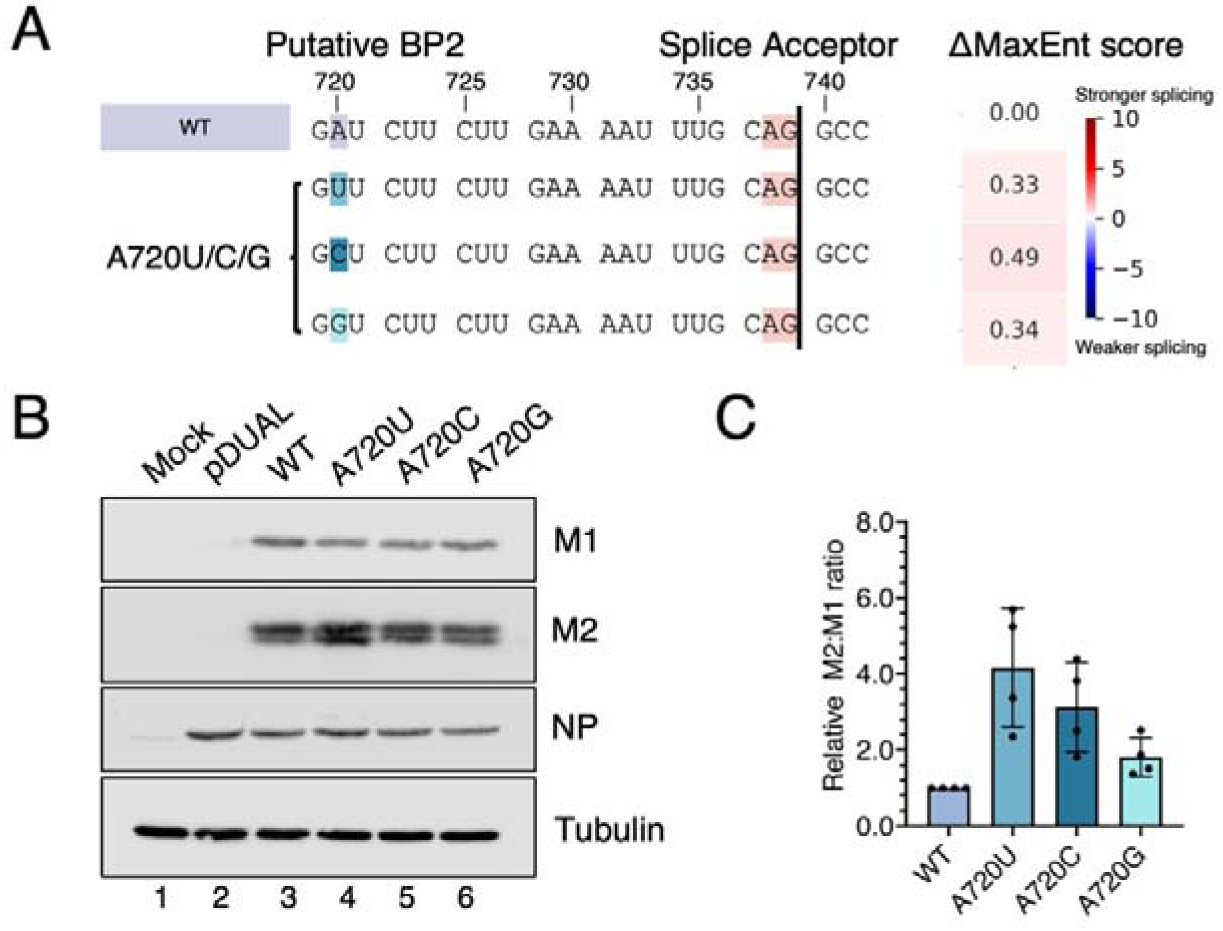
Mutational tests of putative BP2 at nucleotide 720. (A) Sequences of mutants with substitutions at the putative BP. Mutated sites are highlighted in blue and the segment 7 SA is highlighted in orange. Heatmap showing predicted ΔMaxEnt (mutant – WT) scores for indicated SA mutants. Positive values indicate the mutation is predicted to increased SA strength. (B) HEK 293T cells were co-transfected (or mock transfected) with the minireplicon components, NS segment and the indicated segment 7 or pDUAL empty vector plasmids. Cells were lysed and analysed by SDS-PAGE and western blotting as labelled. (C) Densitometric quantification of the relative M2: M1 ratio was analysed by an ordinary One-way ANOVA test with Dunnett’s post-test relative to WT (N=4). Absence of significance symbols = P>0.05. Error bars indicate the standard deviation.

As noted above (Figure 1), another potential BP in segment 7 is predicted at nucleotide 713, in addition to the one at position 720. To test if this adenosine was the BP, we mutated it to a guanine (Figure 5A) and examined segment 7 splicing in the minireplicon system. Unlike the A720G change, A713G reduced both mRNA2 and M2 levels (Figure 5B, C), the latter was statistically significant (Figure 5D). However, splicing was not completely blocked, as expected if A713 was the sole branch point. This suggests that while A713 is important for splicing, it is not the only potential BP residue. Instead, the segment 7 BP could be loosely defined [83], and nearby adenosines could be used in the absence of the primary BP [84]. To test this, we mutated two additional adenosines at nucleotides 714 and 715 (A713-5G). This mutant showed a more pronounced drop in mRNA2 and M2 levels, but low levels of splicing still occurred (Figures 5B, lane 6; 5C lane 7, 5D). However, further addition of the A720G mutation (A713-5G+A720G) led to barely detectable mRNA2 levels and undetectable M2 expression (Figure 5B, lane 7, Figure 5C lane 8, Figure 5D). These results support the hypothesis that segment 7 can use more than one BP adenosine, at positions 713-715 and 720.

**Figure 5.**
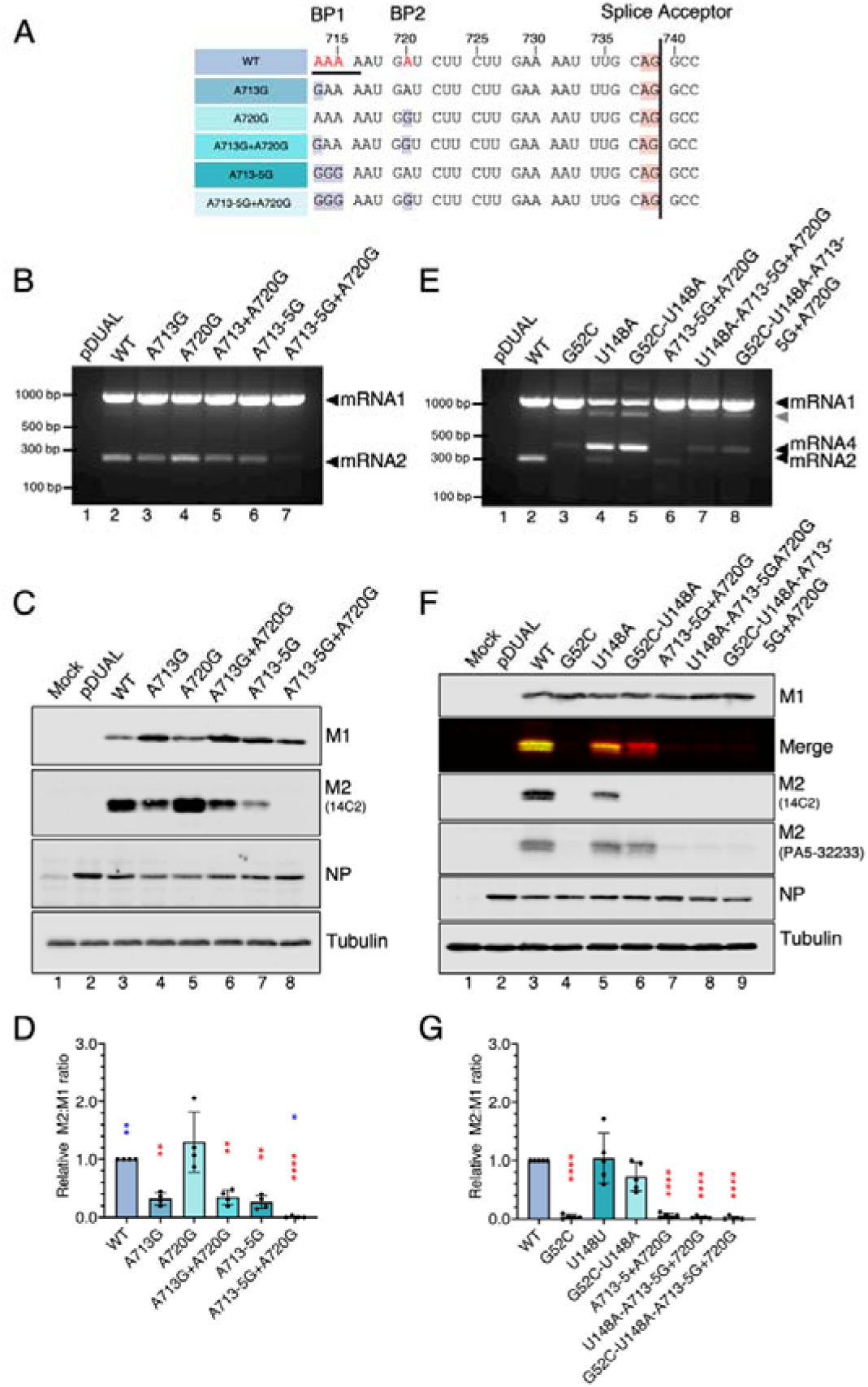
Mutational identification of the segment 7 BP. (A) Mutational strategy for identifying BP adenosines in segment 7. Adenosines that were predicted as potential BPs are shown in red on the WT sequence, mutations are highlighted in purple, while the SA is highlighted in orange. The region of the SRSF5 binding motif overlapping with BP mutations is underlined. (B-G) HEK 293T cells were co-transfected (or mock transfected) with the minireplicon components, NS segment and the indicated segment 7 or pDUAL empty vector plasmids, harvested at 48 h and analysed by (B, E) RT-PCR and agarose gel electrophoresis and (C, F) western blotting as labelled. The grey arrowhead indicates a probable heteroduplex of mRNA1 and 4 PCR products [87]. (D, G) M2:M1 ratios were quantified by densitometry of replicate experiments and normalised to the value of the WT construct. The relative M2:M1 ratios were analysed by an ordinary One-way ANOVA test with Dunnett’s post-test relative to WT (N=4-5). **** = P<0.0001; *** = P<0.001; ** = P<0.01; * = P<0.05; absence of significance symbols = P>0.05. Error bars indicate the standard deviation. Red asterisks indicate statistically significant differences compared with WT, whereas blue asterisks in 5D indicate statistically significant differences compared with A713G.

A previous study identified serine and arginine-rich splicing factor 5 (SRSF5) as a factor that interacts with nucleotides 707-715 to promote mRNA2 splicing [85], overlapping with our putative BPs. To distinguish whether the absence of mRNA2/M2 in the A713-5G+A720G mutant was caused by the loss of BP function or lack of SRSF5 binding, we further tested the mutations in the context of mRNA4 splicing, which does not depend on SRSF5 [85]. Since WT PR8 only produces low levels of mRNA4, we altered the segment splicing pattern to improve mRNA4 and consequently M42 protein expression, by introducing U148A to upregulate the mRNA 4 SD and G52C to destroy the mRNA2 SD [21]. As expected, single mutation of U148A upregulated mRNA4 at the expense of mRNA2, without abolishing its production (Figure 5E, lane 4), while G52C severely reduced mRNA2 levels (Figure 5E, lane 3). The combination of the two mutations reduced mRNA2 below the level of detection (Figure 5E, lane 5). We then added the set of A713-5G-A720 potential BP adenosine mutations onto these two backgrounds. This caused a similar reduction in mRNA4 levels to that seen for mRNA2 (compare lane 6 with 7 and 8). Next, to differentiate accumulation of the M2 and M42 proteins we used two anti-M2 antibodies with differing specificities: the 14C2 antibody recognises residues 4 to 17 of the PR8 M2 protein but does not bind M42 [21,86], while a polyclonal antibody (PA5-32233) that was raised against amino acids 11-97 of M2 recognises both M2 and M42 but is unable to distinguish them. M1 levels were relatively consistent across all segment 7 plasmid transfection samples, while M2/M42 levels showed greater variation. The G52C mutant (disrupted mRNA2 SD) did not produce detectable amounts of M2 as expected, and its M42 protein expression was also below the detection limit (Figure 5F, lane 4). For the U148A mutant, 14C2 reactivity was weaker than the WT, whereas PA5-32233 reactivity was comparable to WT, indicating that U148A produced both M2 and M42 (Figure 5F, lane 5). The double mutant G52C-U148A produced only M42, as evidenced by reactivity with PA5-32233 but not with 14C2 (Figure 5F, lane 6). Consistent with the RT-PCR data, once 713-715 and 720 adenosines were mutated, both M2 and M42 production were dramatically reduced (Figure 5F, lanes 7-9; quantification in 5G). Given that SRSF5 does not affect mRNA4 splicing and M42 expression [85], the much-decreased mRNA4 and M42 production seen here suggests that adenosines 713-715 and 720 play a distinct role in segment 7 splicing beyond serving as an SRSF5 binding motif. Overall, these data suggest that segment 7 has redundancy in BP function to mediate mRNA2 and mRNA4 splicing, although we could not directly identify the specific BP(s) utilised.

### The strength of the segment 7 SD is set by nucleotides C51 and C55

The SD motif of cellular pre-mRNAs is recognised by base-pairing between the U1 snRNP and 9 nucleotides either side of the mRNA splice site (Figure 6A) [31]. The IAV segment 7 mRNA2 SD sequences are highly conserved (except for nucleotide 55) and a close match to cellular consensus. The two exceptions are nucleotides 51 and 55 (Figure 6A), although the presence of a pseudo-uridine in U1 snRNA may allow some flexibility in base pairing for the latter residue. To determine how sequence variation within the mRNA2 SD region influences segment 7 splicing, we investigated all possible polymorphisms *in silico* using MaxEnt to predict the change in splice-site intrinsic strength. As expected, results indicated that the GU dinucleotide at positions 52 and 53 were the most important sites, with all changes drastically reducing SD strength (Figure 6B). Most other possible changes were also negative, or neutral, but substitutions C51G and C55A where the PR8 SD diverges from consensus were predicted to enhance mRNA SD strength. To test these predictions experimentally, we created the panel of all possible point mutations in the mRNA2 SD region and analysed their behaviour in the minireplicon assay as before. RT-PCR analysis of segment 7 mRNAs indicated that altering the three nucleotides in the exon side of the motif had only slight effects on splicing levels, although as predicted, mutation C51G made splicing slightly stronger, visibly changing the mRNA2:mRNA1 ratio (Figure 6C, lane 11). Densitometric analysis of the RT-PCR products indicated a 2.3 ± 0.3-fold (n=3) increase compared to WT. Also consistent with the MaxEnt predictions, all variations of G52, U53, G56 and U57 inhibited mRNA2 splicing, mostly drastically (Figure 6D, lanes 3-8 and Figure 6E, lanes 6-11). Defective recognition of the mRNA2 SD was also associated with upregulation of mRNA4 levels, presumably due to competition between these two SDs for U1 snRNP interaction [21]. Conversely, C55, the only intronic position with any variability in segment 7, was more tolerant of mutation. Consistent with the IAV consensus, C55U showed a similar level of splicing to WT, while mutating C to a purine (especially A) made splicing stronger (mRNA2:mRNA1 ratio 3.1 ± 0.1-fold higher than WT, n = 3). To provide a more quantitative measure of segment 7 gene expression, we used western blotting to measure viral protein expression. Since some of the introduced point mutations are non-synonymous in M1/M2 and amino acid substitutions in the M2 ectodomain interrupt the 14C2 anti-M2 antibody epitope, the polyclonal anti-M2 antibody (PA5-32233) was used. NP and NS1 levels were similar between all samples, indicating equivalent levels of transfection (Figures 6F-H, Figure 6I for quantification). Consistent with the RNA analyses, point mutation C51G reduced M1 accumulation and gave higher levels of M2 (although not statistically significant), consistent with mRNA1 oversplicing. The C55A and C55G substitutions formed premature stop codons in M1 so predictably, no M1 was detected in these two samples (Figure 6H, lanes 5 and 7). Even so, C55A showed increased M2 levels (Figure 6H, lane 5), which when quantified against α-tubulin levels, were significantly higher than WT (Figure 6I). Also consistent with the RT-PCR data, M2 antibody reactivity was absent (or very much reduced; U57 series) from all mutants having mRNA2 splicing defects (protein expression in Figure 6G, lanes 5-12 and 6H, lanes 8-13). However, even though defective mRNA2 SD sequences upregulated mRNA4 SD recognition, M42 levels did not reach the detection limit in our experiments.

**Figure 6.**
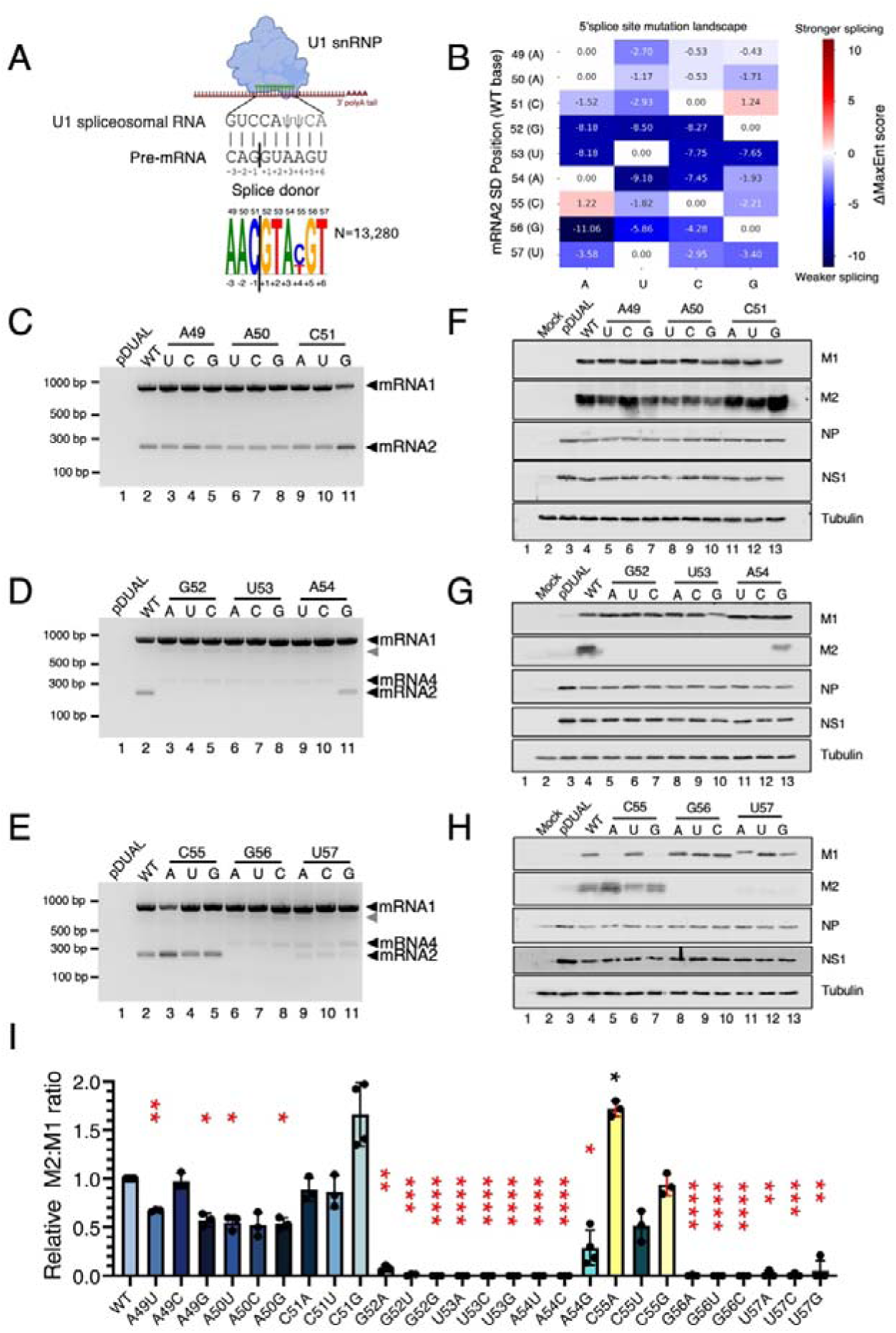
Mutational analysis of the mRNA2 SD sequence. (A) Cartoon diagram of base-pairing between U1 snRNP and mRNA SD sequences. Ψ represents pseudouridine, an isomer of uridine found in U1 snRNA. Sequence logo indicating conservation of the segment 7 mRNA SD site, generated from 13,280 IAV segment 7 sequences. (B) Heatmap showing predicted ΔMaxEnt (mutant – WT) scores for indicated SD mutants. Positive values indicate the mutation is predicted to increased SD strength, whereas negative values indicate a weaker SD. (C-H) The mRNA2 SD was systematically mutated and individual constructs were tested by minireplicon system. HEK 293T cells were co-transfected (or mock transfected) with viral polymerase and NP (3PNP), NS segment and the indicated segment 7 or pDUAL empty vector plasmids. Lysates were harvested 48h post-transfection and (C-E) segment 7 mRNA detected by RT-PCR followed by agarose gel electrophoresis. The grey arrowhead indicates a probable heteroduplex of mRNA1 and 4 PCR products [87]. (F-H) Viral protein expression levels were analysed using western blotting. (I) M2:M1 ratios were quantified by densitometry of replicate experiments and normalised to the value of the WT construct. For mutants C55A and C55G (yellow bars), M2:tubulin ratios were calculated due to the presence of premature stop codons in the M1 ORF, leading to undetectable M1 protein expression. Statistical analysis was performed by ordinary One-way ANOVA tests with Dunnett’s post-test relative to WT (N=3-4). Red asterisks indicate tests for M2:M1 ratios, black asterisk for M2:tubulin. **** = P<0.0001; *** = P<0.001; ** = P<0.01; * = P<0.05; absence of significance symbols = P>0.05. Error bars indicate the standard deviation.

Thus, combining the RT-PCR and western blotting results, all point mutations of nucleotides 52, 53, 54, 56 and 57 inhibited mRNA2 splicing. Only two mutations, C51G and C55A, increased segment 7 splicing. Thus, the mRNA2 SD is strong, but not entirely optimised to support mRNA2 SD recognition and splicing.

### Splicing regulation in viable viruses

So far, we analysed segment 7 mRNA splicing using a minireplicon system to avoid any confounding effects of splice site mutations on the primary amino acid sequence of M1 and segment-specific packaging signals. However, the kinetics and regulation of segment splicing in the minireplicon system may not fully reflect that of the authentic virus. Therefore, to test how the *cis*-acting splicing elements regulate mRNA production in the context of viral infection, as well as to explore the evolutionary conflicts between the multiple functional elements in segment 7, we attempted to rescue a panel of viruses containing mutations that altered the balance of mRNA1 and mRNA2 production in the minireplicon system. For the PPT, we redesigned the mutations to strengthen the PPT by introducing pyrimidine substitutions while avoiding repetitive nucleotides (Figure 7A; 728C-30U, 731-32Y and 728C+730-32Y), to avoid the potential for transcriptional slippage by the viral polymerase [80]. To weaken the PPT, the two pyrimidines immediately upstream of the SA motif were instead substituted with purines (734-36R; the purine at position 736 was swapped from G to A to prevent the creation of an AG dinucleotide), based on the results from a deep mutational scan of the PR8 M1 protein indicating that it can tolerate the corresponding amino acid substitution (L237E, Figure 7A and B) [88]. We also cross-referenced other key mutants that had previously been tested in the minireplicon system for potential effects on M1 function because most of these mutations in the splicing motifs are non-synonymous in M1. (Figure 7B). Some of the sites altered in our study exhibited a high mutational tolerance, for example, amino acid D232 (nucleotide changes A720U, A720C and A720G), and E235 (728-30U and new mutant 731-32Y), suggesting a lower likelihood of impact on M1 function. In contrast, the lower diversity of amino acids recovered at positions 9 - 11 of M1 suggests greater functional conservation, raising the possibility that mutations at these positions could affect M1 function. Therefore, any phenotypes associated with mutations at these positions could reflect effects on splicing, M1 function or both.

**Figure 7.**
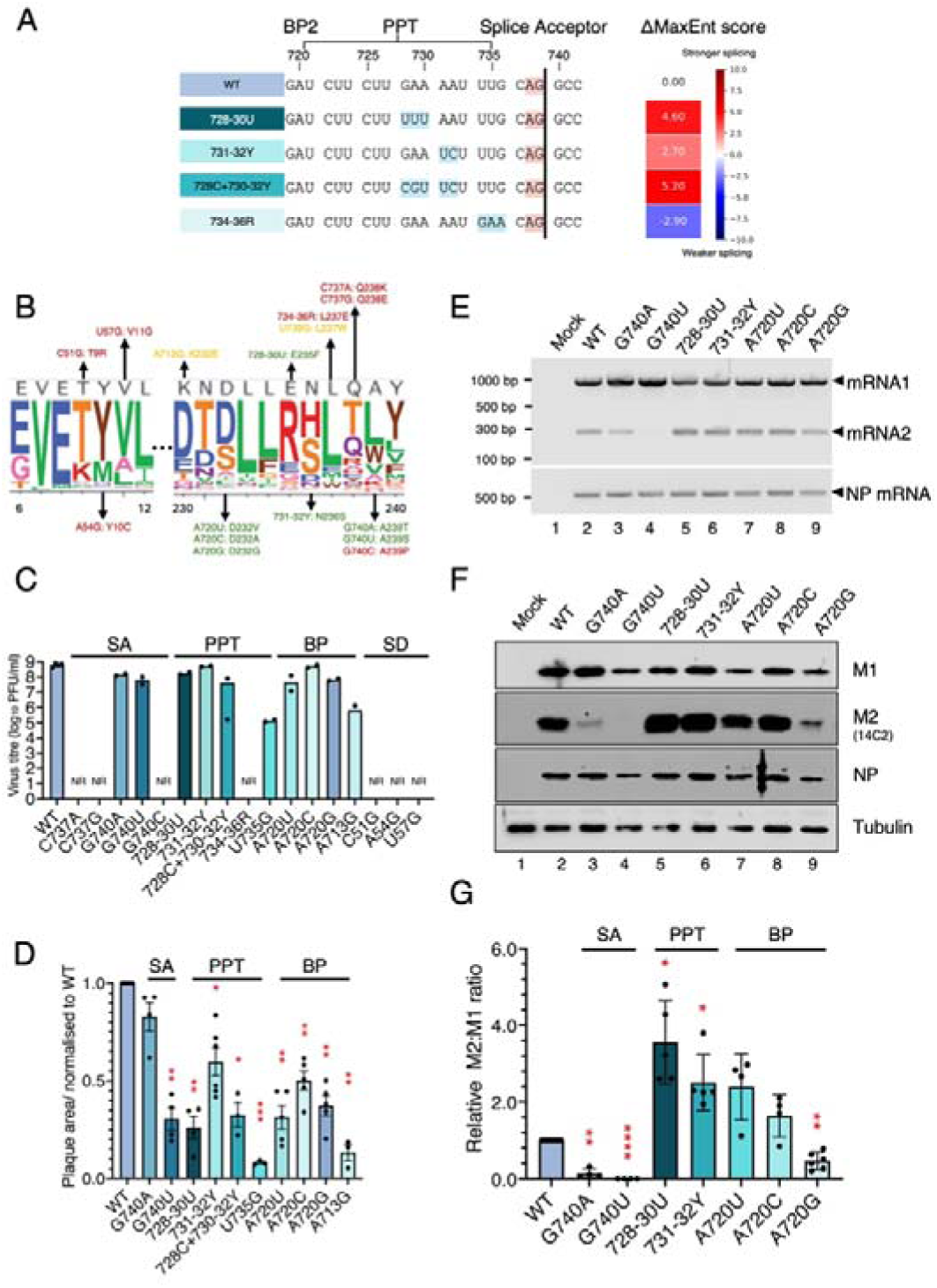
Virus phenotypes following mutation of *cis*-acting segment 7 splice elements. (A) Left - design of a new panel of mutants with mutated PPTs. Mutations are highlighted in blue. The canonical splice acceptor site is shown in orange and the positions of BP2 and the PPT are labeled. Right - heatmap showing predicted ΔMaxEnt (mutant – WT) scores for indicated SA mutants. Positive values indicate the mutation is predicted to increased SA strength, whereas negative values indicate a weaker SA. (B) Logo plot representing the amino acid preferences at each M1 residue site [88]. Mutations tested here are labelled, and amino acid substitutions are shown. Mutants that could not be rescued are shown in red; mutants exhibiting titres less than 10% of WT are coloured in yellow; mutants which grew to near WT levels (sufficient to allow high multiplicity synchronous infections) are coloured in green. (C) Titres of WT and mutant viruses grown from MDCK cells. Data represent the mean of at least two independent rescues. NR – not rescued. (D) Quantification of the plaque sizes of PR8 WT and mutant viruses. Data are the mean ± SEM of 5-20 plaques normalised to the average WT value from at least three independent experiments. (E, F) The effects of splice element mutations on segment 7 gene expression. MDCK cells were infected (or mock-infected) with the indicated viruses at MOI 5 and (E) total RNA isolated at 6 hours post-infection (h p.i) and the indicated mRNAs were detected by RT-PCR, or (F) protein lysates collected at 8 h p.i and viral protein expression analysed by SDS-PAGE and western blotting. (G) M2:M1 ratios were measured by densitometry, normalised to WT and compared using an ordinary One-way ANOVA test with Dunnett’s post-test (N=4-6). **** = P<0.0001; *** = P<0.001; ** = P<0.01; * = P<0.05; absence of significance symbols = P>0.05. Error bars indicate S.D. Panel B was adapted with permission from Hom et al., Journal of Virology 2019;93:e00161-19 (CC BY 4.0 https://creativecommons.org/licenses/by/4.0/).

We next assessed whether the redesigned PPT mutations were predicted to alter splicing in the intended direction. MaxEntScan predicted that pyrimidine addition would strengthen splicing, while reduction would weaken it (Figure 7A, heatmap). Viruses were rescued by transfecting bidirectional reverse genetics plasmids into HEK 293T cells, amplified by one passage in MDCK cells, and plaque titred. WT PR8 grew to an average titre of 6×10^8^ PFU/ml and formed large plaques (Figures 7C, D). No mutants with alterations to the SD motif or the nucleotide preceding the SA site could be successfully rescued. Two of three mutants with altered first nucleotides downstream of the SA in the exon grew to WT-like levels, albeit with small-plaque phenotypes, but G740C could also not be rescued. Most PPT mutants rescued successfully but only 728-30U and 731-32Y gave similar titres to WT, although forming significantly smaller plaques. 728C+ 730-32Y replicated to more than 10-fold lower titres than WT, while 734-36R was apparently non-viable. Mutations at position 720 rescued readily and replicated to titres approaching that of WT virus, with only moderately reduced plaque sizes. The BP1 mutation A713G caused a far more severe growth defect, both for titre and plaque size. Overall, more than half the splice site mutations gave viable viruses with only moderate fitness penalties, indicating a degree of evolutionary flexibility in the 3’ splice site sequence motifs.

Next, we investigated segment 7 mRNA splicing levels in cells infected with the set of WT and mutant viruses that grew to sufficient titres to permit high multiplicity infections. Examination of NP mRNA levels by RT-PCR confirmed that all infections had proceeded successfully (Figure 7E). In cells infected with WT PR8, a mixture of unspliced mRNA1 and spliced mRNA2 was detected (Figure 7E, lane 2). Mutations at the nucleotide immediately following the AG (especially G740U) resulted in reduced mRNA2 production, confirming that this position serves as an essential *cis*-acting element for regulating splicing (Figure 7E, lanes 3 and 4). In contrast, mutant viruses containing additional pyrimidines in the PPT enhanced splicing efficiency, shifting the mRNA1 and mRNA2 ratio in viable viruses. The A720G potential BP mutant had slightly lower levels of NP mRNA, perhaps indicating impaired infectivity. However, mRNA2 was still detectable in all the A720 mutant viruses. This result was consistent with the previous observations in the minireplicon system, indicating that nucleotide 720 is either not the BP or not the only nucleotide that can provide that function for segment 7 splicing. Thus overall, the RT-PCR results for segment 7 splicing were consistent with our previous minireplicon assay results. When we analysed segment 7 protein expression from the mutant viruses by western blotting, NP and tubulin levels were comparable in most samples, but M2 levels were far more variable. The G740A and G740U mutations flanking the AG acceptor dinucleotide gave very low and undetectable amounts of M2 respectively (Figure 7F, lanes 3 and 4). Quantification of replicate experiments confirmed that the M2:M1 ratio of these two mutants was significantly lower than that of WT, indicating M2 protein expression defects (Figure 7G). Conversely, the 728-30U and 731-32Y PPT up-mutations led to significantly increased accumulation of M2 protein (Figure 7F, lanes 5-6, Figure 7G). The BP2 mutations at position 720 gave conflicting phenotypes. A720U and A720C matched the minireplicon and viral mRNA results, producing abundant quantities of M2 and a small (but not statistically significant) increase in M2:M1 ratios. However, in the context of virus, A720G drastically underproduced M2, despite only a slight decrease in mRNA2 levels. Nevertheless, M2 expression in all potential BP mutants confirm that A720 is not the sole segment 7 BP. Furthermore, with the exception of A720G, all splicing mutations behaved similarly in virus to minireplicon system, supporting the hypothesis that the core segment 7 splice site motifs are involved in setting the balance of M1 and M2 expression.

Since segment 7 splicing efficiency can be cell-type dependent [19] and the cell types used in our previous experiments (MDCK and HEK293T cells) do not represent the natural target cells of IAV, we tested whether the phenotypic effects of mutating core splice site motifs were conserved in a human respiratory epithelial context. Human lung A549 cells were infected with the panel of segment 7 viruses and segment 7 mRNA splicing was investigated using RT-PCR. The detection of comparable levels of NP mRNA confirmed successful infection (Supplementary Figure 4A). WT and mutant viruses exhibited splicing patterns in A549 cells that were broadly consistent with those observed previously in HEK293T and MDCK cells. Mutations G740A and G740U reduced mRNA2 production (Supplementary Figure 4A, lanes 3 and 4), whereas 728–30U and 731–32Y enhanced splicing (lanes 5 and 6), while all A720 mutants retained detectable mRNA2 expression (lanes 7-9). Viral protein expression correlated with transcript abundance. Compared with the WT virus, G740A and G740U mutants expressed reduced or undetectable levels of M2 protein respectively (Supplementary Figure 4B, lanes 2-4, quantification in 4C). Conversely, higher M2:M1 ratios (albeit not statistically significant) were observed in 728-30U and 731-32Y-virus infected A549 cells (lanes 5 and 6). The A720 viral mutants displayed protein expression patterns broadly consistent with their splicing phenotypes. Interestingly, the M2:M1 ratio for the A720G virus was similar to that of WT virus in A549 cells, leaving the reduced splicing of this virus in MDCK cells as the only evidence for potential cell type- or species-specific differences.

### Conservation of segment 7 splicing regulation beyond PR8

The work described above demonstrated that the PR8 segment 7 core *cis*-acting splice elements play a substantial role in regulating segment 7 mRNA alternative splicing thus the balance of M1 and M2. However, PR8 is a laboratory-adapted strain, so the question arose as to whether this would apply to other strains of IAV. To address this, we introduced representative mutations affecting each core *cis*-acting splice element into the 2009 pandemic H1N1 strain, Cal04. Unlike PR8, this Cal04 strain has limited laboratory-adaptation, with no mutations reported in segment 7 [60]. First, all Cal04 mutants were tested by minireplicon assay in both HEK 293T and A549 cells under the same experimental conditions as used for PR8. When segment 7 splicing was measured using RT-PCR, the WT Cal04 segment produced a mix of unspliced mRNA1 and spliced mRNA2 (Figure 8A, E). Substitutions surrounding the SA AG dinucleotide inhibited splicing, with mutations on the exon side exhibiting lesser effects. Mutations that increased PPT pyrimidine content enhanced splicing whereas purine substitutions reduced it (Figure 8B, F). Spliced mRNA2 was detected in all potential BP mutants but mutation of A713G had the greatest effect (Figure 8C, G), similarly to in PR8. Likewise, the SD mutation C51G increased splicing, while A54G and U57G mutations strongly downregulated mRNA2 (Figure 8D, H). Unlike in PR8, downregulation of mRNA2 production did not increase mRNA4 production in the Cal04 background, most likely because of its weaker mRNA4 SD sequence (MaxEnt score -9.47 versus -1.72). To investigate how splicing alterations influence Cal04 virus fitness, we rescued the set of mutants, amplified them by one passage in MDCK cells and determined virus titres (Figure 8I). Like PR8, Cal04 did not tolerate mutations of the SD or preceding nucleotide of the SA site. The fitness reductions observed in PR8 G740A and G740U were also recapitulated in the Cal04 strain. G740A reached WT-level titres but showed a small-plaque phenotype, whereas G740U exhibited more severe defects (Figure 8I, J). The potential BP1 mutant A713G, which was severely attenuated in PR8, could not be rescued in the Cal04 background, while all BP2 A720 mutants gave WT-level titres. However, mutations that enhanced PPT function were better tolerated in Cal04 than in PR8. Although the 728-30U, 731-32Y and 728+730-32Y mutants formed smaller plaques, all produced comparable titres to WT. Conversely, all PPT-disrupting mutants were non-viable. Taken together, the overall effects of mutations in the core *cis*-acting elements were generally consistent in Cal04 and PR8, supporting the hypothesis that the core *cis*-acting splice elements regulate segment 7 splicing across distinct influenza A virus strains.

**Figure 8.**
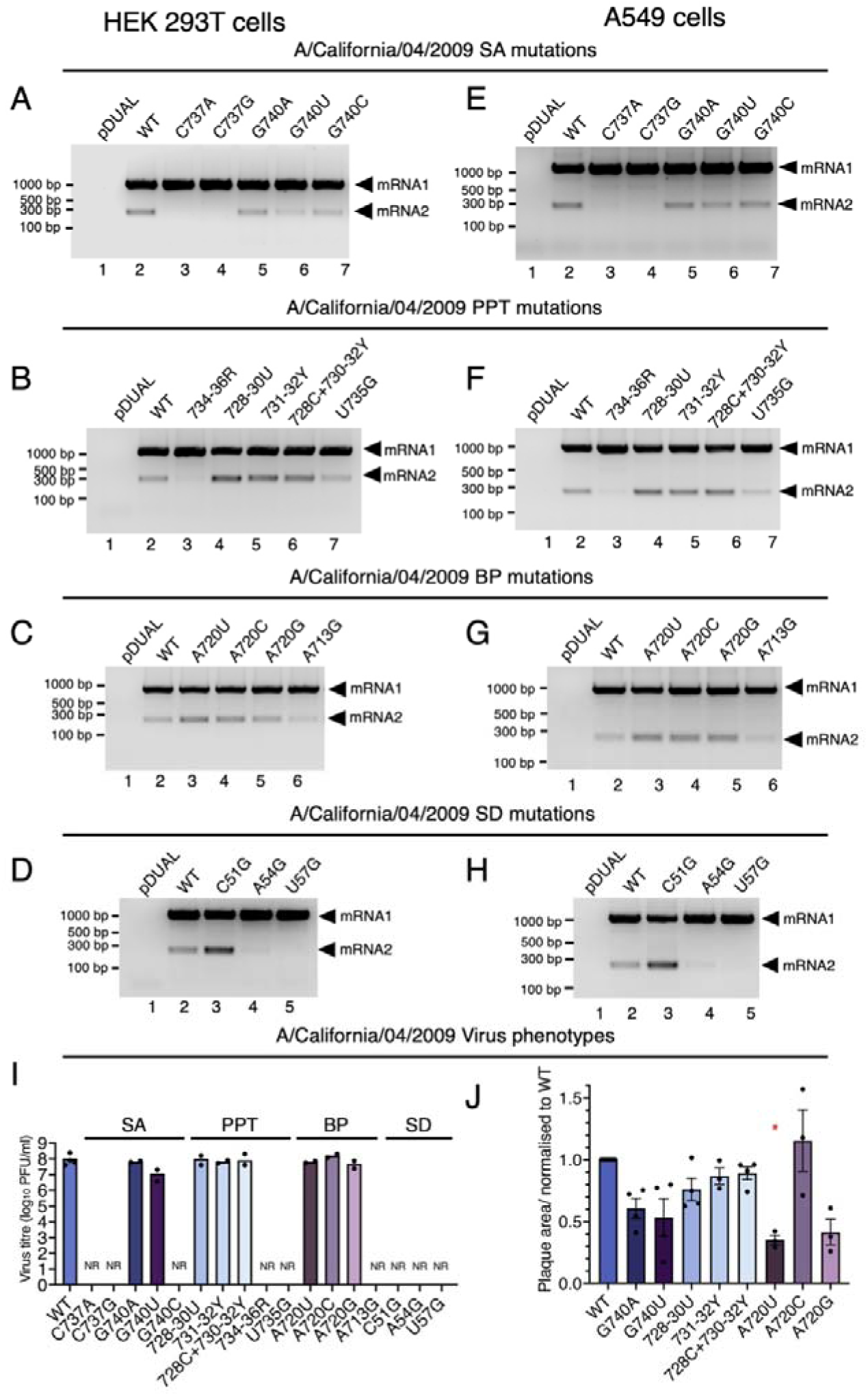
Validation of the effects of core segment 7 splice site mutations in the Cal04 pandemic H1N1 strain. Key mutations in core splice elements were introduced into the Cal04 segment 7. HEK293T (A-D) or A549 (E-H) were co-transfected with the Cal04 minireplicon components, NS segment and the indicated segment 7 or pDUAL empty vector plasmids, harvested at 48h and analysed by RT-PCR and agarose gel electrophoresis. (I) WT and mutant Cal04 viruses were rescued in HEK 293T cells, and propagated in MDCK cells and virus titres were plotted. P1 stock titres are plotted. Dots represent independent rescues and bars indicate the mean. NR – not rescued. (J) Quantification of the plaque sizes of WT and mutant Cal04 viruses. Data are the mean ± SEM of 5-20 plaques normalised to the average WT value from at least three independent experiments.

To investigate whether the segment 7 *cis*-acting splicing elements are conserved across IAV strains from different host species, sequence logos for human, avian, swine, equine, canine, and feline IAVs were generated. The core SD and SA motifs were highly conserved across all host species, with only nt 55 and nt 740 exhibiting single-nucleotide polymorphisms (Supplementary Figure 5). The PPT region from nts 721-735 showed a similarly conserved pattern; although positions 721, 724 and 733 varied between C or T, the five purines at nts 728-732 and the 3’end of PPT remained conserved. Therefore, the core *cis*-acting splice signals of segment 7, including the negative regulatory elements in SD and PPT, are highly conserved among influenza A viruses from diverse host species. While this will reflect selective pressure from the coding sequences, it is consistent with a role in regulating segment 7 mRNA splicing despite potential differences in the host splicing environment.

## Discussion

IAV uses alternative splicing of segment 7 to encode two proteins, M1 and M2, and impaired production of either of these two proteins can reduce virus fitness [21,89-92]. The regulation of segment 7 alternative splicing is complex, and previous studies have reported various control mechanisms, involving *cis*-acting sequences in segment 7 and *trans*-acting protein factors, both host and viral. For example, a polypyrimidine and polycytosine tract downstream of the mRNA2 SD has been shown to influence splicing by binding to the NS1-BP and hnRNP K proteins [36]. Alternative splicing factor 1/splicing factor 2 (ASF/SF2) which binds to a purine-rich stretch downstream of the SA is also involved in mRNA2 splicing as a *trans*-acting element [44]. Potential viral *trans*-acting factors include the viral polymerase binding to the cRNA 5’-promoter element downstream of the 5’-cap-snatched leader sequence to block usage of the mRNA3 SD site, itself proposed to regulate usage of the mRNA2 SD [39]. NS1 also potentially modulates splicing through direct or indirect interactions with mRNA1 and mRNA2 as well as by promoting nuclear export of unspliced mRNA1 [37,93-95]. There may be elements of virus strain-specificity to some of these mechanisms, given (for instance) the variable amounts of mRNA3 production and disagreements over whether or not M2 is essential for virus replication [21,91]. Analyses of the role of NS1 in controlling viral mRNA splicing have also given variable results [38].

Here, we describe how suboptimal splice signals in IAV segment 7 maintain a ratio of spliced and unspliced transcripts and consequently balanced production of M1 and M2 proteins (Figure 9). A strong SA site and potentially redundant BP adenosines promote efficient mRNA2 splicing and M2 protein production. Negative sequence elements in the SD and PPT regions prevent oversplicing to preserve sufficient essential viral protein M1 production from unspliced mRNA.

**Figure 9.**
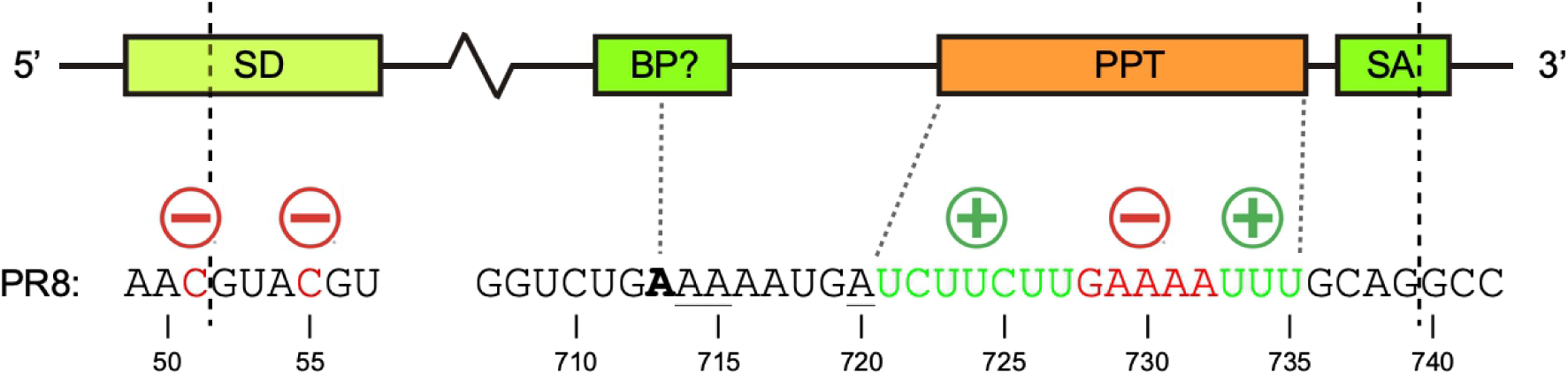
Diagrammatic summary of how IAV segment 7 uses suboptimal splicing signals to maintain the balance of mRNA1/M1 and mRNA2/M2 production. Segment 7 has a strong splice acceptor (SA), a strong branch point (BP; the most likely lariat-forming adenosine indicated in bold with potential alternative adenosines underlined) and a medium-strong splice donor (SD) which allow the virus to produce sufficient M2 protein. A purine-rich stretch in a medium strength PPT prevents oversplicing, thereby preserving levels of unspliced mRNA1 required for the synthesis of the essential M1 protein.

Thus, although multiple factors can regulate segment 7 mRNA splicing, we propose that the four required *cis*-acting elements, SD, BP, PPT, SA are the primary determinants that regulate viral mRNA splicing to a level consistent with virus viability. This model is consistent with the high conservation of the segment 7 splice motifs (Figure 1B, Supplementary Figure 5). It would also maintain a basal regulation of M1/M2 production both across the frequent reassortment events seen in avian and other influenza A viruses that could pair the segment with variants of the viral polymerase and/or NS1 with altered splicing regulatory activity. Given the evolutionary conservation of the basal splicing machinery across birds and mammals [96,97], segment-autonomous regulation may also help underpin a broad host range, as host jump events might place the virus in a cellular environment with altered sets of SRE factors. Further "fine-tuning" of splicing ratios could then evolve for the particular gene constellation and cellular environments for an individual virus lineage or host.

Achieving the correct balance of M1 and M2 exerts a selection pressure on the virus to evolve suboptimal splicing signals. Further sequence constraints arise from maintaining function of the translation products encoded by the mRNAs, as well as any other *cis*-acting functions of the segment. All splice site elements considered in this study overlap with the M1 open reading frame (ORF) and some of them overlap the M2 ORF. Consequently, most mutations that alter splicing efficiency are non-synonymous in M1 and sometimes in M2. The mRNA2 SD sequence is also involved in the segment-specific packaging signal [58]. Mutations in this region therefore might not only change splicing efficiency [30] but also affect virus genome packaging, or even M1/M2 translation via codon bias effects. Consistent with a severe constraint on the segment sequence, all our attempts to rescue three mRNA2 SD mutants failed. A previous study also reported failure to rescue mRNA2 SD mutants, finding that virus could only be generated successfully when the mutations within mRNA2 SD motif (G52 and U53, M1 codons 9 and 10) were synonymous [15]. Other mutants which also failed to produce mRNA2 but which also altered codons 9 or 10 of M1 were non-viable. Therefore, either M1 protein sequence and/or packaging signal sequences are critical in this region, hampering modifications to the splice site. Assessment of changes to codon and codon-pair biases for the PPT mutants did not show a clear correlation with virus fitness (data not shown), and experiment investigation of this mechanism is outside the scope of this study.

In contrast to the relative mutational inflexibility of the SD sequence, the purine-rich stretch (728-732 of segment 7) nested within the PPT was much more tolerant of changes. Introducing pyrimidines which are non-synonymous in M1 to this region produced viable viruses with only limited fitness defects, indicating mutational flexibility. This is consistent with the output of the deep mutational scan of PR8 M1 protein where M1 E235 and N236 sites exhibited a high mutational tolerance for amino acid substitutions [88] and (Figure 7D). Furthermore, this region does not overlap with any currently defined packaging signals in segment 7 [98,99]. Thus, unlike the SD region, which plays multiple essential roles (packaging signal, splice site and encoding M1 and M2) during the viral life cycle, the PPT region has "only" two defined functions: M1 protein coding and acting as a PPT. The combination of relative functional simplicity and high mutational tolerance suggests that the virus can more easily regulate splicing by changing the PPT sequence during evolution. This might explain why the PPT is the most important down-regulator of splicing as it allows more flexibility without severe impacts on viral fitness.

The transfection-based minireplicon assay allows us to investigate the effects of a broad range of mutations on segment 7 splicing. In contrast, infection experiments necessarily focused on mutants that could be rescued and propagated to sufficiently high titres for high-MOI synchronous infection. Thus, the viable PPT mutants examined here represent splicing perturbations that remain compatible with productive virus replication. Our data further suggest that IAV is tolerant of moderate reductions in M1 expression resulting from increased segment 7 splicing. Mutants with a stronger PPT (728-30U and 731-32Y) had approximately two-to four-fold increases in M2:M1 ratios yet still replicated to near-WT titres in the PR8 background. The corresponding Cal04 segment 7 constructs also showed increased levels of splicing and yet in the virus background, replicated to WT levels.

Although differences in replication kinetics are not necessarily captured by endpoint titres measured at 48 or 72 h p.i., these mutants generally exhibited small-plaque phenotypes, indicating slower replication rates and modest fitness penalties. This is consistent with a suggestion that virus budding needs a threshold amount of M1 to start [90]. In this interpretation, M1 abundance in the over-spliced mutants remains above the minimum level to support virion assembly despite altered M1: M2 ratios. These findings further indicate that a single fixed M1:M2 ratio is not needed for virus viability.

In most situations, M2 is considered essential for normal virus replication [21,91]. However, our data show that the PR8 virus can tolerate a low M2 expression level and still replicate to high titres. The PR8 G740U mutant exhibited a severe splicing defect and failed to produce detectable amounts of mRNA2/M2 during high multiplicity synchronous infection. Despite this, it only exhibited modest growth defects: a slightly reduced titre compared to WT PR8 and a small-plaque phenotype. This finding aligns with previous studies [15,21,58,91,92] showing various IAV strains can undergo multiple cycles of replication despite lacking M2 ion channel activity or overall M2 expression. However, the degree of replication attenuation seen varies by over 1000-fold, depending on strain and mutational strategy, raising the possibility that the requirement for M2 may vary across different strains.

To avoid confounding effects from mutations affecting M1 or M2 protein function and/or packaging signals, we based much of our study around a non-infectious minireplicon system, which may not completely recapitulate the temporal aspects of virus infection [14,89,100]. Nevertheless, when we compared our results from minireplicon and viable virus infection, the majority of mutants behaved similarly. The exception was the A720G mutation in the PR8 virus background in MDCK cells where splicing was weaker, in contrast to PR8 virus in A549 cells and PR8 and Cal04 minireplicons performed in 293T cells or Cal04 minireplicons in A549 cells, where splicing was similar to WT (Figures 4B, 5B, 8C and 8G). Presumably this reflects cell type differences, although additional experiments would be required to test this hypothesis.

## Supplementary data

Supplementary data is available at NAR online.

## Data Availability

The data underlying this article are available in Open Science Framework repository at https://osf.io/uxp2y/overview?view_only=273fdaffd8f143d4852934f3efd81e52.

## Funding

RMP, ERG and PD acknowledge Institute Strategic Programme grant support from the UK Biotechnology and Biological Sciences Research Council (grant no. BBS/E/RL/230002C).

## Conflict of interest

The authors declare no competing interests.

## Acknowledgements

We thank Professor Daniel Perez for supplying the Cal04 reverse genetics system, as well as Professor Chris Smith and Dr Colin Sharp for helpful discussions. The graphical abstract was created in BioRender. Zhou, S. (2026) https://BioRender.com/0osc6de. We acknowledge the use of ChatGPT to assist with debugging of scripts for classifying host species for IAV segment 7 sequences. The authors reviewed, tested, and modified all AI-assisted code as necessary.

## Author Contributions

SZ: conceptualisation, data curation, formal analysis, investigation, methodology, visualisation, writing (original draft), writing (review & editing).

RMP: Funding acquisition, methodology, visualisation, writing (review & editing)

HWC: Data curation, investigation, software

HML: Methodology, writing (review & editing)

EG: Methodology, supervision, writing (review & editing)

PD: Conceptualisation, data curation, formal analysis, funding acquisition, project administration, resources, supervision, writing (original draft), writing (review & editing)

## Notes

### Competing Interest Statement

The authors have declared no competing interest.

### Summary of Updates

Various revisions to the text and figures after two rounds of helpful comments from reviewers

