## Supplementary information for "Identification of *cis*-Acting Elements that Regulate Influenza A Virus Segment 7 Differential mRNA Splicing"

**Supplementary Data: Zhou et al.**

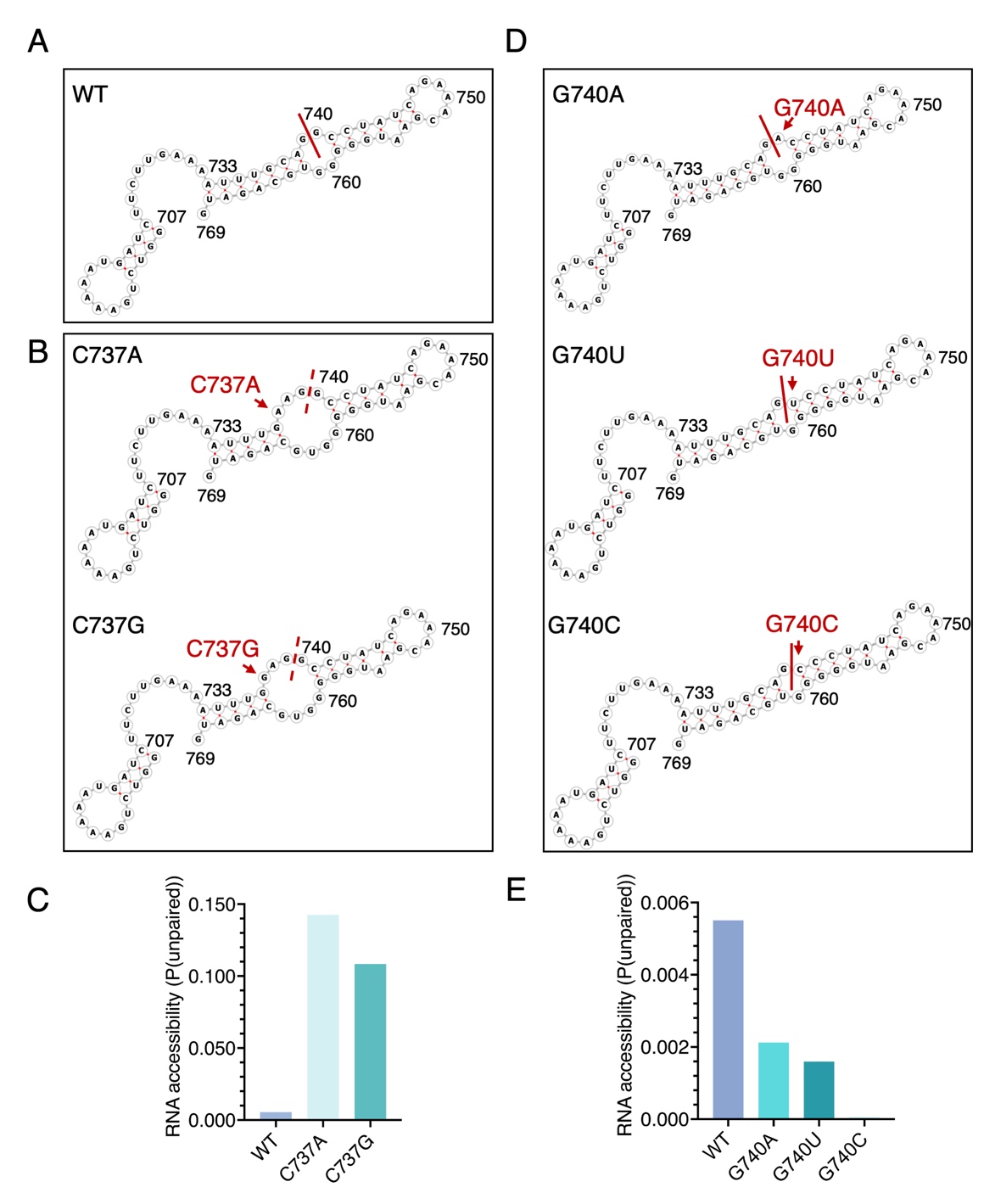

**Supplementary Figure 1. Predicted RNA secondary structures and the RNA accessibility at the segment 7 SA region.** (A, B and D) RNAfold-predicted WT and mutant RNA secondary structures. The splice site is indicated by a red line or dashed line. The splice site is indicated by a solid line when splicing was detected in the minireplicon assay, and the by a dashed line when splicing was not detected. (C and E) RNAplfold estimates of local RNA structural accessibility of C737 mutants (C) and G740 mutants (E). The mean probabilities that the tetranucleotide CAG|G is unpaired are shown.

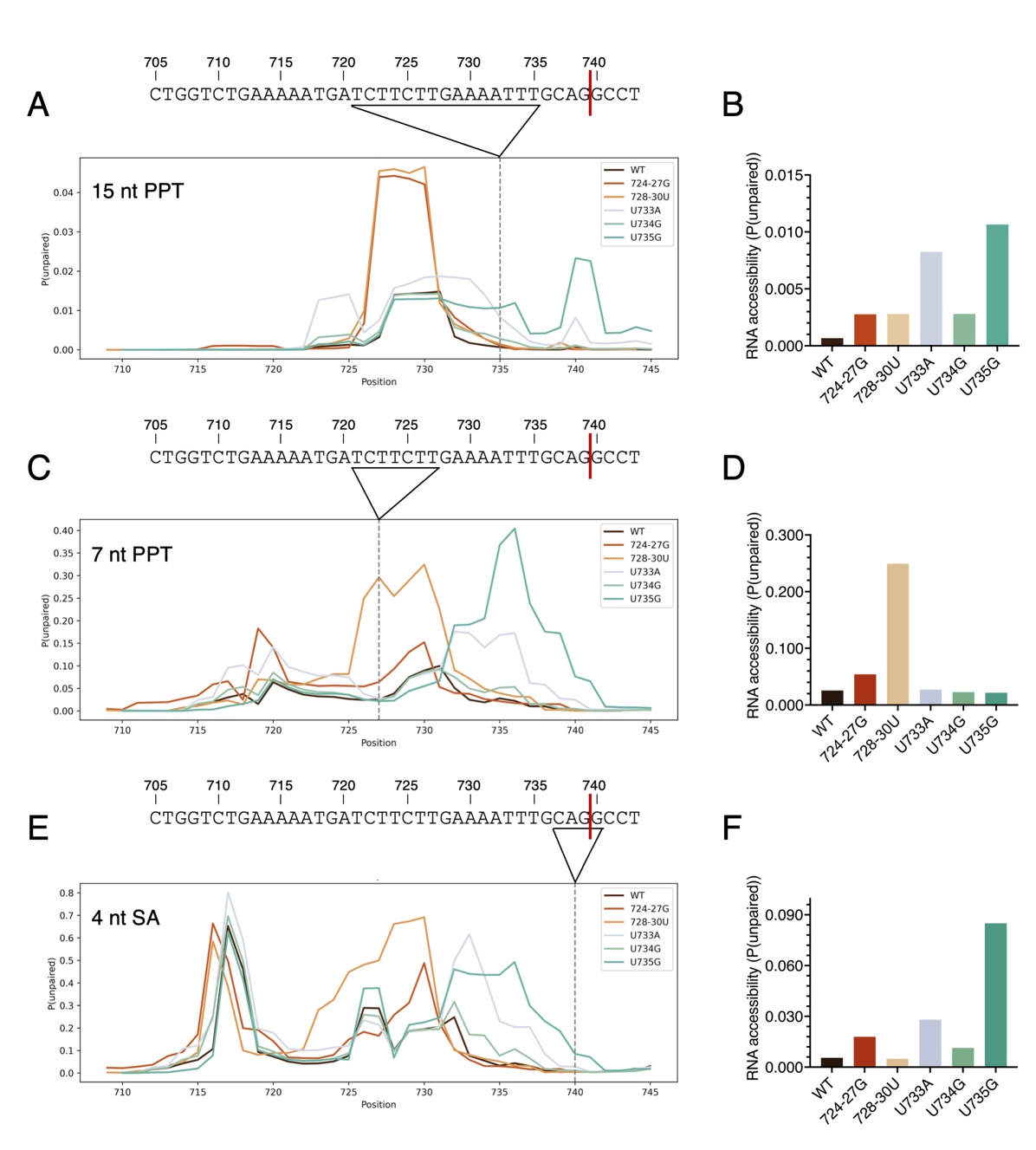

**Supplementary Figure 2. RNA accessibility analysis of PPT mutants.** Local RNA accessibility of (A, B) the putative 15 nt PPT identified in this study (nt 721–735), (C, D) the previously predicted 7 nt PPT by Moss *et al*. (46, 54) (nt 721–727), and (E, F) the splice acceptor (SA) motif (nt 737-740) were calculated using RNAplfold. Line plots show the mean probability that stretches of 15, 7, and 4 consecutive nucleotides, respectively, ending at the plotted nucleotide position are unpaired. Dashed lines mark the 3′ end of the sequence element under analysis. The probabilities that these positions are unpaired are presented separately for WT and mutant RNAs in panels B, D, and F.

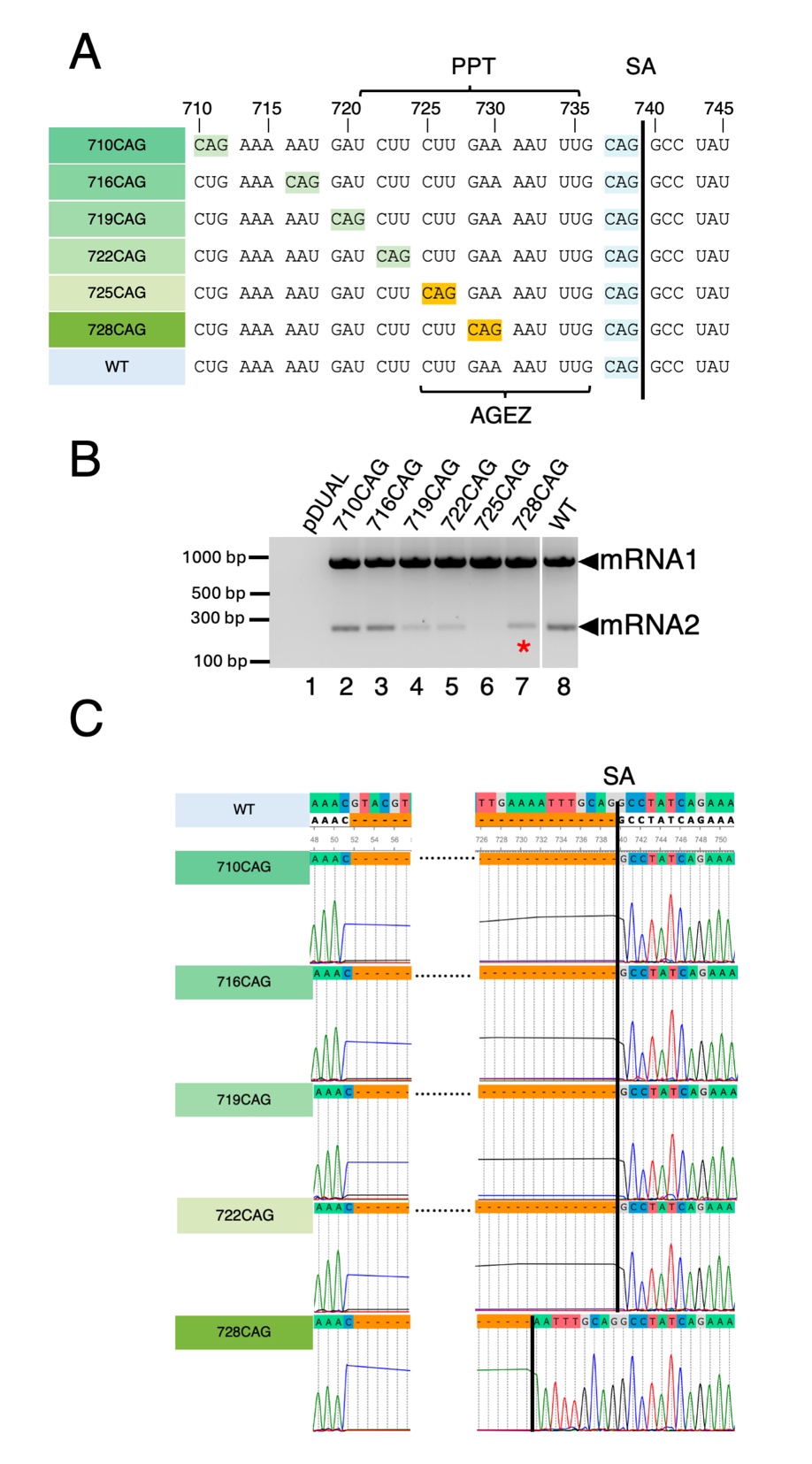

**Supplementary Figure 3.** **Mapping the segment 7 AGEZ.** (A) Design of mutants with novel CAG motifs upstream of the canonical SA site. CAGs that are ignored are highlighted in green and ones that interfere with use of the canonical SA are highlighted in amber. The canonical SA is highlighted in blue. (B) HEK 293T cells were co-transfected (or mock transfected) with the mini-replicon components, NS segment and the indicated segment 7 or pDUAL empty vector plasmids, then harvested at 48 h and analysed by RT-PCR and agarose gel electrophoresis. The red asterisk indicates a spliced mRNA2 band arising from usage of the novel AG, unlabelled ones are normal mRNA2 produced by canonical SA. (C) Sanger sequencing results of PCR products corresponding to the indicated mutant spliced transcripts (mRNA2). The SA sites are indicated by black lines.

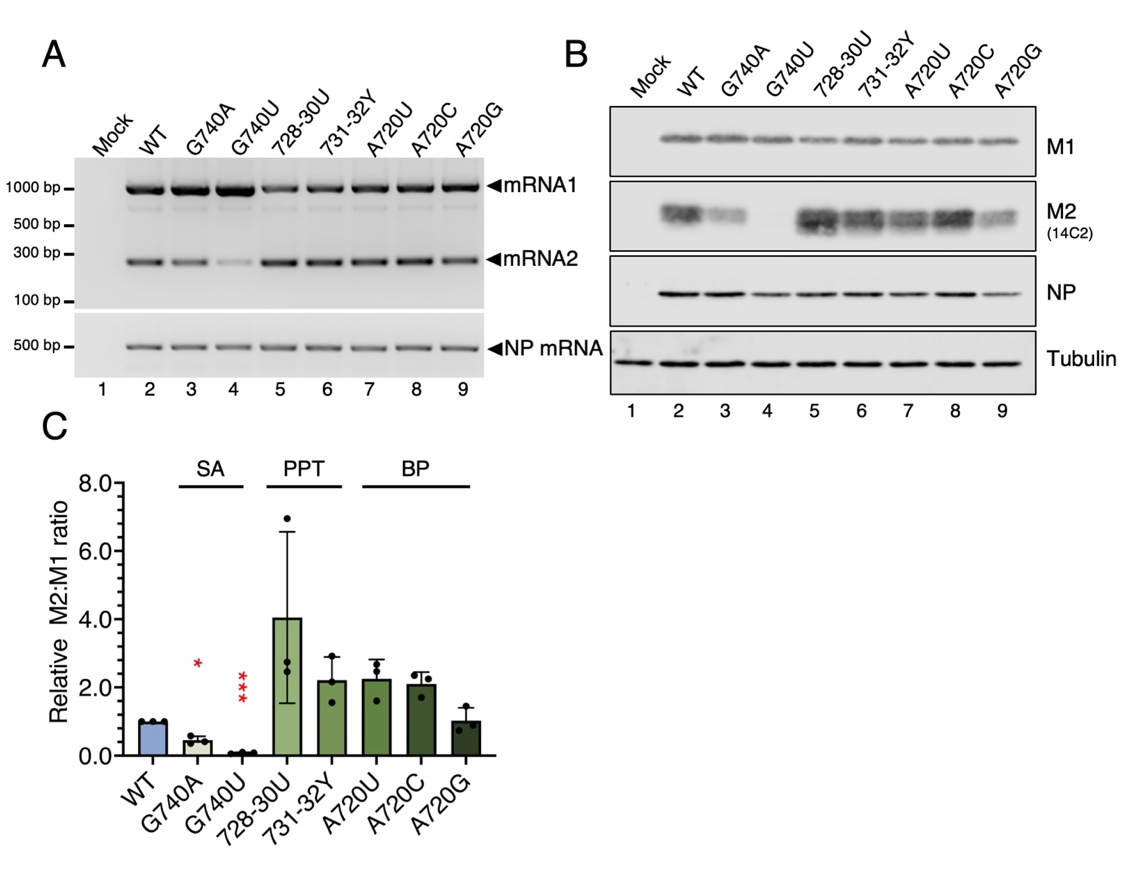

**Supplementary Figure 4. The effects of segment 7 splice site mutations on viral mRNA splicing and protein expression in human lung epithelial cells.** (A, B) A549 cells were infected (or mock-infected) with the indicated viruses at an MOI of 5. (A) Total RNA was isolated at 6 h p.i and the indicated mRNAs were detected by RT-PCR. (B) Cells were lysed at 8 h p.i. and viral protein accumulation analysed by SDS-PAGE and western blotting. (C) M2: M1 ratios were measured by densitometry, normalised to WT and compared using an ordinary One-way ANOVA test with Dunnett’s post-test (N=3). **** = P<0.0001; *** = P<0.001; ** = P<0.01; * = P<0.05; absence of significance symbols = P>0.05. Error bars indicate the standard deviation.

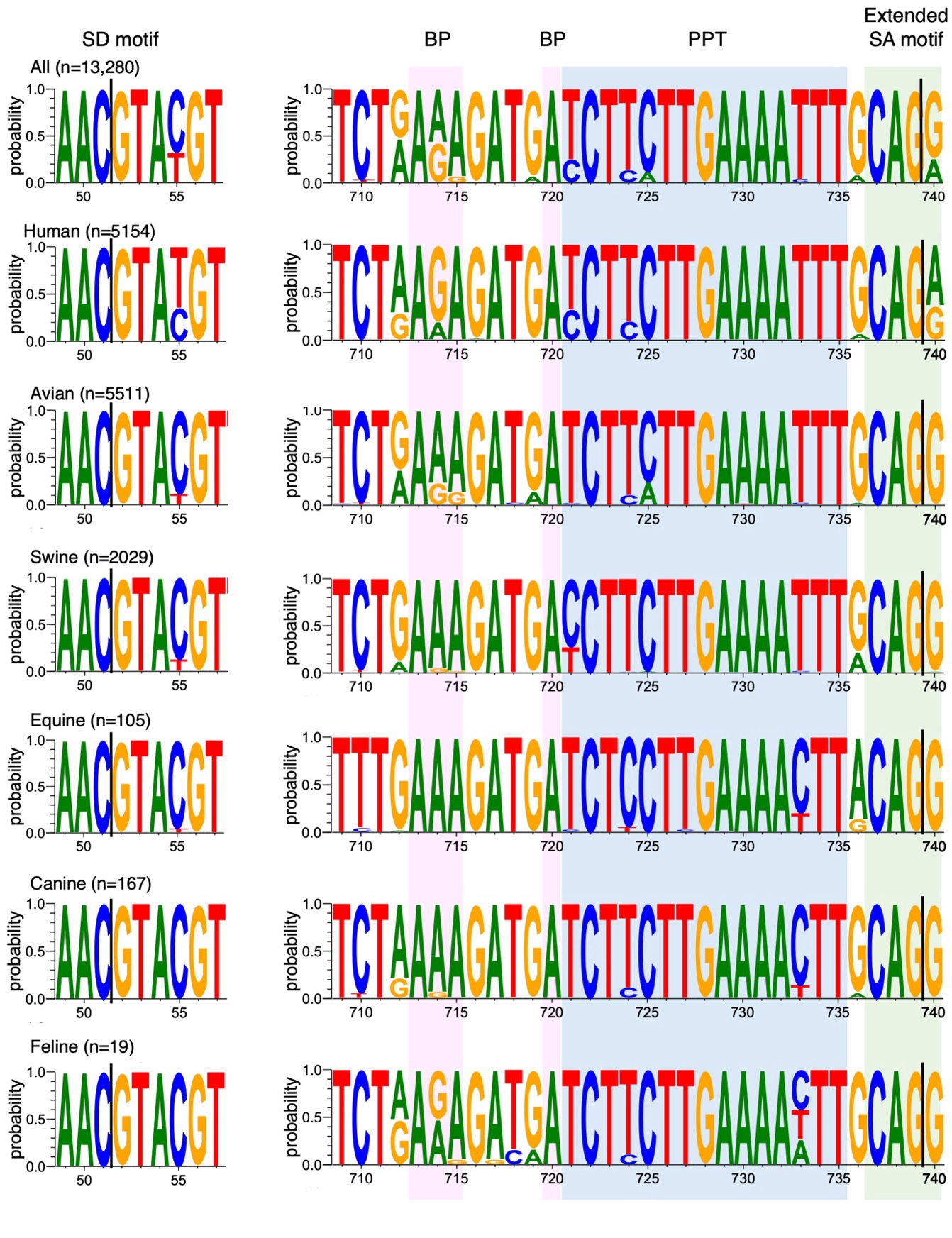

**Supplementary Figure 5. Sequence logos of IAV segment 7 core *cis*-acting splice signals broken down by host type.** SD, BP, PPT and SA motifs of IAV segment 7 from viruses isolated from the indicated species/orders were used to generate sequence logos. Splice sites are indicated by black lines.

**All citations in the Supplementary Data file use the numbering in the main text bibliography**
